# Extensive nuclear introgression accompanied ancient mitochondrial capture in the common ancestor of two sister hare species

**DOI:** 10.64898/2026.09.14.750963

**Authors:** José Costa, João Pedro Marques, Liliana Farelo, Eugénio Silva, Sofia Granja Martins, Angela Trentacoste, Greger Larson, Joel Alves, João Queirós, Paulo Célio Alves, Pierre Boursot, José Melo-Ferreira

## Abstract

Pleistocene climatic oscillations promoted contact and hybridization among hare species, leaving lasting genomic signatures. Two sister species, *L. castroviejoi* (broom hare, endemic to Iberia) and *L. corsicanus* (Italian hare, endemic to Italy), carry fixed mitogenomes from *L. timidus* (mountain hare), an Arctic species widespread in southern Europe before deglaciation. Whether such turnover was accompanied by nuclear introgression, and how it affects their genomes today, remains unclear. Here, we date the ancient mitochondrial replacement in the broom-Italian hare ancestor using whole mitogenomes incorporating Iron Age samples. Coalescent modelling of complete genomes infers a substantial mountain hare contribution (24%) to this ancestor ∼900 Kya (∼5N generations). Accordingly, phylogenetic segmentation identified genomic segments consistent with mountain-to-broom/Italian hare introgression, covering ∼17.8% of the autosomes, close to ∼18% expected under drift, given admixture’s timing and magnitude. Genome polarisation corroborates this estimate, with 25–28% of the mountain hare genome polarising with broom and Italian hares rather than its sister Iberian hare, decreasing to 15–18% at diagnostic sites. The introgression segments concentrate fixed-introgression sites and show sharp transitions of ancestry at their borders. Highly introgressed regions are gene-dense, and regions of reduced introgression are repeat-rich, suggesting that recombination shapes the introgression landscape. They also harbour genes with mitochondrial function that mirror candidates linked to mito-nuclear coadaptation in the Iberian hare, hinting that mitonuclear interactions may influence which variants persist. Our study illustrates how combining demographic, phylogenomic and polarisation-based methods can quantify and localize extensive nuclear introgression, even in the absence of non-introgressed reference populations.

## Introduction

Introgression, once thought to be a rare process in nature, is a crucial evolutionary mechanism. The integration of foreign alleles via hybridization and subsequent introgression, can significantly shape genetic variation (Hedrick 2013; Tigano and Friesen 2016; Seixas et al. 2018; Rosser et al. 2024). Numerous studies across a wide range of biological systems have shown that introgression can accompany speciation leaving behind lasting genomic signatures (Reich et al. 2010; Rheindt and Edwards 2011; Lamichhaney et al. 2015; Ferreira et al. 2021; Qian et al. 2023). Furthermore, such introgressed variation has often been reported to provide adaptive benefits, namely by facilitating rapid adaptation to novel environments (Huerta-Sánchez et al. 2014; Liu et al. 2015; Qian et al. 2023; North et al. 2024). Characterizing the genomic consequences of introgression, including its extent and distribution across the genome, is therefore essential to understand how introgressed variation contributes to species evolution.

The most frequently reported instances of introgression concern the mitochondrial genome (Toews and Brelsford 2012). Although this pattern may partly reflect ascertainment bias, it warrants particular attention because mitochondria play a central role in key cellular processes, including energy production, metabolism, and apoptosis. As such, the mitochondrial genome harbours considerable adaptive potential (Gemmell et al. 2004; Da Fonseca et al. 2008; San-Millán 2023). Mitochondria interact closely with numerous nuclear-encoded genes, and incompatibilities between mitochondrial and nuclear genomes have been documented in several systems (Gemmell et al. 2004; Smith et al. 2010; Hill 2017; Sloan et al. 2017; Barreto et al. 2018). Additionally, the “mother’s curse”, i.e. the accumulation of mitochondrial mutations that are neutral or beneficial in females but deleterious in males, can further exacerbate these incompatibilities if not counteracted by compensatory nuclear mutations (Gemmell et al. 2004; Smith et al. 2010; Seixas et al. 2018). Despite the constraints on mitochondrial introgression, studies have demonstrated compensatory responses in the nuclear genome, suggesting that selection can favour nuclear changes to restore mito-nuclear compatibility (Barreto et al. 2018; Hill 2020; Zhao et al. 2025). Consequently, it is important to assess whether cases of extensive mitochondrial introgression are accompanied by introgression of interacting nuclear genes and to understand how such events shape patterns of variation across the nuclear genome.

The Iberian Peninsula harbours three hare species that provide a rich system for addressing this question: the broom hare (*Lepus castroviejoi*), the Iberian hare (*L. granatensis*), and the European hare (*L. europaeus*), hereafter referred to as the brown hare for clarity. All three species have experienced repeated episodes of extensive introgression of mitochondrial genomes from the same origin, the mountain hare (*L. timidus*) (Melo-Ferreira et al. 2007; Seixas et al. 2018; Souto et al. 2025). The best-studied case to date is the Iberian hare, which exhibits across the Iberian Peninsula a south–north gradient of mitochondrial introgression (Seixas et al. 2018). In contrast, nuclear introgression does not generally follow this geographic pattern, instead appearing more homogeneously distributed across the Peninsula and typically at low levels (Seixas et al. 2018). Seixas et al. (2018) proposed a demographic and biogeographic explanation for this discrepancy. Nevertheless, they also identified outlier nuclear genomic regions that mirror the geographic pattern of mitochondrial introgression, including genes involved in mitochondrial metabolism. However, the extent to which such patterns reflect broader, non-random distributions of introgressed variation across the genome remains unclear.

The broom hare provides a more straightforward system to address this question, as mitochondrial introgression is complete: this species shows no native mitochondrial haplotypes and instead carries exclusively mountain hare–like haplotypes (Melo-Ferreira et al. 2012; Ferreira et al. 2021). This marked discrepancy between the nuclear and mitochondrial genomes has been validated through coalescent simulations (Melo-Ferreira et al. 2012). Such a pattern raises the expectation that coevolved nuclear genomic regions may also have undergone extensive introgression. Its sister species, the Italian hare (*L. corsicanus*), currently allopatric, likewise exhibits complete mitochondrial introgression and is therefore crucial for a comparative framework (Melo-Ferreira et al. 2012).

Previous studies have investigated nuclear introgression in these sister species and reported very low levels of standing admixture from any other hare species from Europe (<1%) (Souto et al. 2025). This contrasts sharply with their mitochondrial genomes, which appear to be entirely of mountain hare origin. However, because these analyses treated each recipient species as a parental reference for the other, they were only able to detect differential introgression occurring after the divergence of the broom and Italian hares. Consequently, introgression shared by both species through admixture into their common ancestor would remain largely undetected.

A first objective of this study is to determine whether mitochondrial DNA replacement occurred independently in the two hare sister species or only once in their common ancestor, and to date this event using a combination of modern and Iron Age mitochondrial genomes. We also ask to what extent large-scale mitochondrial introgression was accompanied by nuclear introgression, how introgression is distributed along the genome, and whether similar genomic regions are involved in the Iberian hare system affected by introgression of the same mitochondrial lineage. Using a combination of analyses exploiting different aspects of our whole genome resequencing data, we infer that the mitochondrial replacement occurred during an ancient major admixture event of the mountain hare with the ancestor of the broom and Italian hares, that also left extensive traces in their autosomal genomes, including a significant portion of fixed mountain ancestry.

## Methods

### Sampling, laboratory procedures, sequencing and bioinformatic processing

We analysed whole-genome sequences from 22 broom hares, 25 Italian hares (20 mainland and 5 from the introduced population in Corsica), 6 mountain hares, 10 Iberian hares, 10 brown hares, and 8 snowshoe hares, including both newly generated (37 genomes) and previously published data (44 genomes; Seixas et al. 2018; Giska et al. 2019; Souto et al. 2025) (Fig. 1; Table S1). Samples were obtained from the CIBIO-InBIO collection and provided by collaborators or collected during permitted hunting seasons. For newly sequenced specimens, genomic DNA was extracted from ear or internal organ tissue using the EasySpin Genomic DNA Tissue Kit (Citomed, Lisbon, Portugal), with samples preserved in ethanol or RNAlater. Aiming at high-coverage sequencing, TruSeq DNA PCR-Free libraries (Illumina) with ∼350 bp insert size were prepared and dual-indexed. Additional libraries were prepared using double-indexed libraries following (Meyer and Kircher 2010), aimed at subsequent low-coverage sequencing. All libraries were sequenced as 150 bp paired-end reads on an Illumina NovaSeq 6000 platform (Novogene, Cambridge, UK).

**Figure 1.**
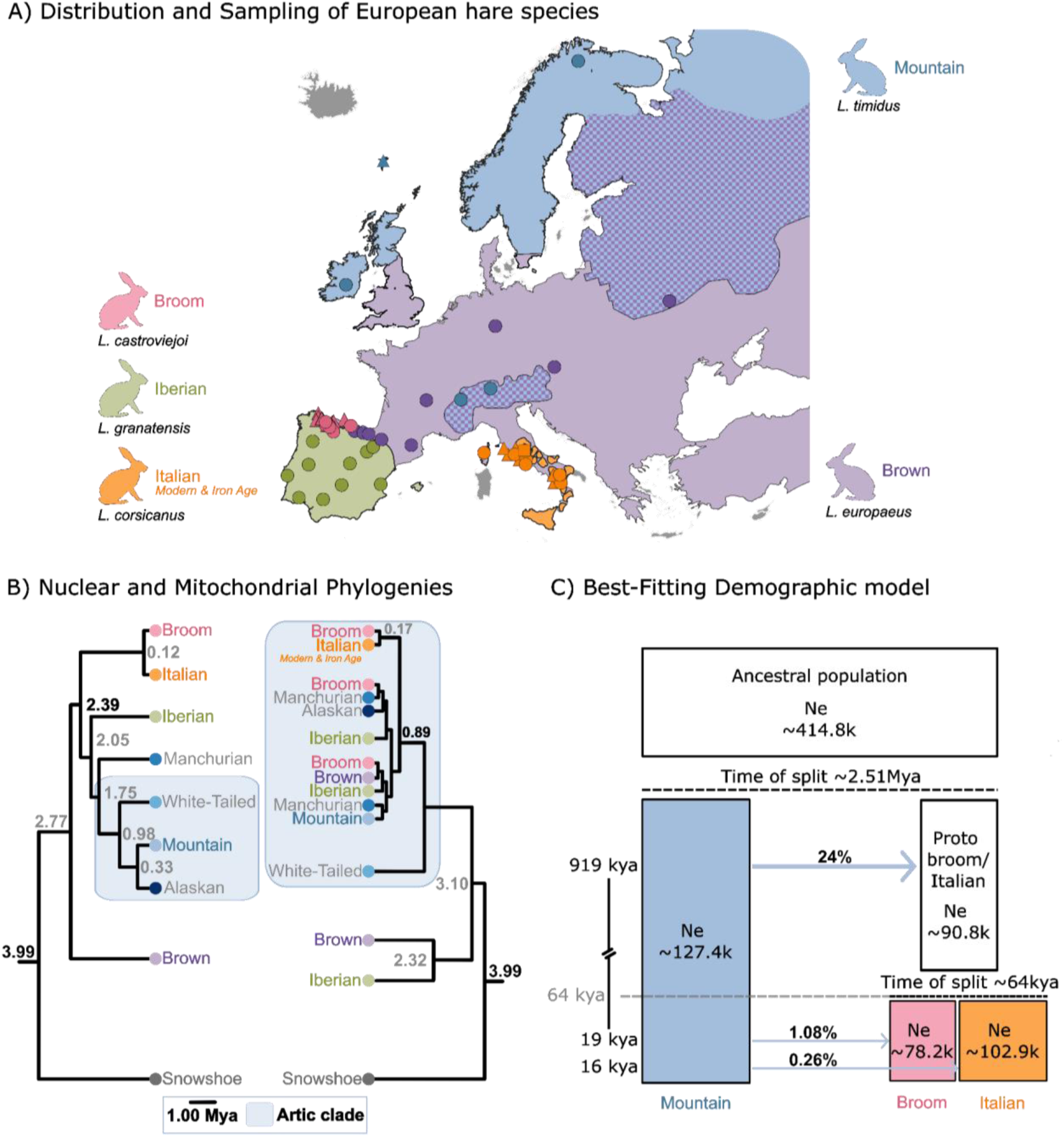
Sampling locations and phylogenomic relationships. **A)** Geographic distribution of sampled individuals and species from Europe. Samples from the Far East Russia (mountain hare; N=1) and North America (snowshoe hare; N=8) are not shown in the map. Species ranges are shown by coloured shading; dot colours indicate species’ identities, and symbol shape denotes sequencing depth (circles: high coverage; triangles: low coverage; squares: Iron Age). **B)** Nuclear species phylogeny (left) adapted from Ferreira et al. (2021), with topology and node ages, and mitochondrial DNA phylogeny (right) adapted from Costa et al. (2024) with divergence times inferred in this study. Node labels indicate divergence times (Mya: million years ago); grey species names denote taxa not sampled in this study. **C)** Best-fitting BPP model of divergence and gene flow among mountain, broom and Italian hares, showing three introgression events. Blue arrows indicate introgression direction; percentages show estimated introgression proportions, and coloured boxes represent extant species. **Alt text:** Sampling locations, nuclear and mitochondrial phylogenomic relationships, and graphical representation of the demographic model.

Adapters and terminal bases were trimmed and reads shorter than 36 bp were removed using Trimmomatic v0.40 (Bolger et al. 2014). Reads were mapped to the brown hare reference genome (Michell et al. 2024) using bwa-mem2 v2.2.1 (Vasimuddin et al. 2019) and processed with samtools v1.8 (Li et al. 2009). Duplicate reads were removed with Picard, and indels were realigned using GATK v3.2-2 (Auwera and O’Connor 2020). SNP calling was performed per species using bcftools mpileup v1.10.2 (Li 2011), after which VCFs were merged and indels removed.

Three Italian hare bone samples were used for ancient mitochondrial DNA analysis. This material originated from Cavità 254, a stone quarry at Orvieto (Italy) that was rapidly infilled during the late Iron Age (Etruscan period, Trentacoste 2021). The hare bones were not directly radiocarbon dated. The chronology of the deposit was based on direct AMS dating of an associated chicken bone from the same well-sealed stratigraphic context (775–541 BC, 95% confidence; Best et al. 2022, George et al. 2017). Of seven hare bone samples initially screened for ancient DNA, three (hereafter Lcor_Iron_125, Lcor_Iron_127 and Lcor_Iron_129) were selected for mitochondrial targeted enrichment based on their endogenous DNA content. Laboratory work followed standard ancient DNA protocols at a dedicated facility (Palaeogenomics & Bio-Archaeology Research Network, University of Oxford), and sequencing reads were processed and mapped following established ancient DNA pipelines (details in Supplementary Text S1). The three consensus mitochondrial genomes had mean coverages of 53.4×, 63.3×, and 141.7×, with 3.0%, 2.4%, and 2.5% missing data, respectively.

### Population structure

Genotype likelihoods were estimated from genomic data using ANGSD v0.935 under the samtools model (Li 2011), to account for variation in sequencing depth. Analyses were restricted to high-quality data (minimum base and mapping quality ≥30), retaining only uniquely mapped reads. SNPs were identified using a likelihood ratio test (p ≤ 1×10⁻⁶) and filtered for a minimum minor allele frequency (MAF) of 0.034 for the broom hare and 0.037 for the Italian hare, with thresholds determined by the number of haplotypes, and for presence in at least 50% individuals. Major and minor alleles were inferred from allele frequencies, with ancestral states defined using the brown hare reference genome. To limit biases from uneven coverage, maximum depth thresholds were set to (2 × coverage mode × number of samples) (132 for broom hare; 80 for Italian hare). Site allele frequency likelihoods (SAF) and per-site read counts were computed for downstream analyses.

Population structure was inferred using NGSadmix v33 (Skotte et al. 2013) for K = 1–5. The optimal K was determined based on log-likelihoods and the Evanno method (Evanno et al. 2005), implemented in CLUMPAK (Kopelman et al. 2015). For the selected K, individual ancestry proportions were visualized geographically using QGIS.

### Mitochondrial phylogeny reconstruction

Contemporary mitochondrial genomes were assembled and annotated using MitoCatch [available at: https://github.com/evochange/MitoCatch], a Snakemake-based pipeline applied to filtered FASTQ files. Reads were subsampled per individual and assembled with Trinity v2.9.1 (Grabherr et al. 2011), followed by annotation using MITOS2 v2.1.7 (Donath et al. 2019). Read subsets are detailed in Table S2. MitoCatch was developed in this study, with documentation and source code available at [https://github.com/evochange/MitoCatch]. Using this approach, complete mitochondrial genomes were recovered for 72 samples (88%) (Table S2).

For samples lacking complete mitochondrial assemblies, a reference-based approach was applied using NOVOPlasty v4.3.5 (Dierckxsens et al. 2016), recovering complete mitochondrial genomes for 4 of 9 missing cases (Table S2). Assemblies were generated with default parameters using species-specific cytochrome *b* sequences, obtained from successful MitoCatch runs, as seeds. The mitochondrial genomes of the three Italian hare ancient samples, obtained from Iron Age archaeological remains at Orvieto, Italy (radiocarbon dated to 776– 540 BC via an associated faunal bone from the same stratigraphic context; Best et al. 2022) were then added to the mitogenome dataset.

The 13 mitochondrial protein-coding genes and two rRNA genes were concatenated and aligned using MAFFT v7.505 (Katoh and Standley 2013). A maximum-likelihood phylogeny was then inferred in IQ-TREE v3.1.3 (Wong et al. 2026) under a GTR-Gamma model. This tree was subsequently time-calibrated using least-squares dating, as implemented in LSD2 (To et al. 2016). The snowshoe–brown hare most recent common ancestor (MRCA) was constrained to 3.99 Mya (95% HPD: 3.59–4.36 Mya; Ferreira et al., 2021), and the three ancient samples were set as non-contemporaneous tips, dated to 775-541 BC (Best et al. 2022) and converted to years before present. Confidence intervals were obtained from 100 resampling replicates. As a confirmatory analysis, the same divergence times were also independently estimated in BEAST v2.7.8 (Bouckaert et al. 2019) using only the contemporary samples and a calibrated Yule prior (further details in Supplementary Text S2). The resulting tree was visualized in FigTree v1.4.4 (Rambaut 2018).

To provide broader phylogenetic context, we incorporated the mtDNA phylogeny of Costa et al. (2024), which was reconstructed from cytochrome *b* gene fragments and included additional taxa not represented in our dataset. Divergence times estimated in the present study were attributed to the corresponding shared nodes of this expanded topology, allowing visualization of a more comprehensive phylogeny, while retaining the temporal estimates derived from our complete mitogenome analyses.

### Demographic modelling

A total of 1,000 intergenic loci (1 kb each), separated by at least 50 kb, were randomly selected across autosomes in numbers proportional to chromosome length. This design targeted putatively neutral, independent regions while ensuring genome-wide representation and avoiding overrepresentation of larger autosomes.

Multilocus sequence alignments were generated from BCF files after filtering for sites with no missing data, a minimum quality score ≥30, and sequencing depth between 5× and 224×. Alignments were built using five high-coverage individuals per species (Lcas1, Lcas2, Lcas6, Lcas7, Lcas9, Lcor6-Lcor10, Ltim1-Ltim4, Ltim6) for the broom, Italian, and mountain hares. FASTA files were produced using bcftools consensus, with IUPAC (International Union of Pure and Applied Chemistry) ambiguity codes assigned to heterozygous sites and Ns to missing data. These loci were used to test for gene flow among the three target species using the Bayesian method implemented in BPP v4.8.0 (Flouri et al. 2018). The Jukes-Cantor (JC) substitution model was used to compute site likelihoods. A gamma prior was assigned to the root age (τ = 2, 2000), while node ages were assigned a uniform Dirichlet prior. Population size parameters were also assigned a gamma prior (θ prior = 2, 2000). MCMC step lengths were automatically tuned (finetune = 1), and samples were recorded every 1,000 iterations.

Gene flow was modelled using the MSC-I (Flouri et al. 2020) and MSC-M (Flouri et al. 2023) frameworks. In the former model, we allowed for one to three migration events from the mountain hare into the ancestor of the broom and Italian hares, as well as into each derived lineage independently. A multispecies coalescent model without gene flow (MSC) (Flouri et al. 2018) was also included for comparison. In MSC-I models, introgression probabilities were assigned a Beta (1,1) prior, whereas MSC-M models used a Gamma (20,1) prior on migration rates. MCMC chains were run for 500,000 iterations (burn-in 40,000; sampling every 2 steps), with extended runs of 1,000,000 iterations for MSC-I models with three migration events to ensure convergence. Models were compared using the Akaike Information Criterion (AIC).

### Genome segmentation according to local phylogeny

We then looked for introgressed genomic segments based on genetic distances between samples of different species using Saguaro (v0.1; Zamani et al. 2013). This method uses a Hidden Markov Model coupled with neural networks to partition the genome into groups of segments sharing similar pairwise distance patterns (called cacti) among samples. The analysis was run sequentially with 1–13 cacti to explore alternative local phylogenetic configurations across the genome, using a subset of individuals and biallelic sites.

The dataset comprised 20 high-coverage individuals: five broom hares (Lcas1–Lcas5), five Italian hares (Lcor6–Lcor10), two Iberian hares (Lgra1, Lgra10), three mountain hares (Ltim1, Ltim2, Ltim4), two brown hares (Leur7, Leur10), and one snowshoe hare (Lame1) as outgroup (Table S1). Individuals were selected to maximise genetic diversity while minimising sample size and thus calculation time.

The VCF file was filtered to retain biallelic sites with a minimum quality score of 30, less than 40% missing data, a maximum sequencing depth of 224, and a minor allele frequency (MAF) ≥ 0.09. The filtered dataset was converted to the binary format required by Saguaro using VCF2HMMFeature, and distance matrices were visualised as unrooted trees in R. Cacti with a phylogeny compatible with introgression of mountain into broom/Italian hares, and discordant from the known species phylogeny, were retained as candidates for introgression (cacti 4 and 12 were identified; see Results).

### Circular bootstrap resampling

To test whether genomic segments of cacti of interest identified by Saguaro were associated with genomic features, we generated replicates of the segments of interest using a circular bootstrap procedure implemented in a custom Python script (https://github.com/evochange/circular-bootstrap). This approach was inspired by similar methods described by Yassin et al. (2016), Nouhaud et al. (2022) and Ebdon et al. (2024). For each set of segments representing a Saguaro cactus, 2,000 replicates were generated by randomising fragment positions while preserving fragment sizes and inter-fragment distances. Chromosomes were treated as circular, allowing fragments to wrap around chromosome boundaries and thereby maintaining the spatial structure of the original regions. The empirical value of the statistic of interest was then compared to the distribution of the values among the bootstrap replicates.

### Basic statistics

Absolute genetic divergence (Dxy) between mountain and either broom or mainland Italian hare samples was estimated using a custom Python script, based exclusively on high-coverage individuals. Sites were required to have a minimum quality score of 30, less than 40% missing data, and a maximum coverage of 224 reads.

D-statistics were calculated using Dsuite Dtrios v0.5 (Malinsky et al. 2021), which estimates D and f4-ratio statistics for all possible species trios. Analyses were performed on the same dataset described above, excluding monomorphic sites, using default settings and the -- abbaclustering option. This option tests whether ABBA sites are significantly clustered along the genome, as expected under introgression, rather than occurring as isolated sites generated by homoplasy or substitution-rate variation. The resulting clustering statistics provides a significance test for introgression that is robust to such confounding processes.

### Genome polarisation

Genomic patterns identified by Saguaro were further evaluated using the genome-wide polarisation framework implemented in *diem* v1.0.1 (Baird et al. 2023), through its Python implementation diempy (Setter et al. 2026). Analyses included five high-coverage individuals each of the broom hare (Lcas1, Lcas6-Lcas8, Lcas9), Italian hare (Lcor6–Lcor10), mountain hare (Ltim1–Ltim4, Ltim6), Iberian hare (Lgra1, Lgra4–Lgra6, Lgra10), and brown hare (Leur2, Leur3, Leur5, Leur6, Leur10). The brown hare samples were masked and thus did not participate in the polarisation process. Diagnostic index (DI) thresholds were determined using the “touchdown DI” procedure suggested in diempy, whereby the DI threshold is progressively adjusted until individuals on either side of the barrier approach hybrid index (HI) values of 0 and 1.

### Comparison of gene introgression patterns with the Iberian system

We compiled a list of 99 genes identified by Seixas et al. (2018) as candidates of interest in the context of mountain hare introgression into the Iberian hare. This included 88 genes showing introgression at high frequencies and 11 mitochondrial-associated genes whose geographic patterns of introgression paralleled those of mitochondrial DNA in Iberia. To assess whether these candidate genes were preferentially represented within regions of high introgression in the broom and Italian hares (i.e., belonging to cacti of interest in the Saguaro analysis) we calculated the ratio between the total base-pair coverage of candidate genes and the total base-pair coverage of all genes overlapping these regions. This empirical ratio was then compared with a null distribution generated from circular bootstrap replicates (see above).

### Functional enrichment

Enrichment of candidate Saguaro cactus regions for genes and coding sequences (CDS) was assessed by comparing the number of base pairs overlapping gene and CDS annotations in the empirical cactus regions with the corresponding distributions generated from the circular bootstrap replicates.

The 0.1% largest cactus 4 and 12 fragments and the 0.1% largest genomic regions lacking these cactus fragments were examined for the presence of genes of interest. Gene Ontology (GO) enrichment analyses were then performed for the set of genes overlapping these regions using g:Profiler (Kolberg et al. 2023), with the European rabbit genome as reference. The background set comprised all annotated genes in the regions analysed by Saguaro.

The 0.1% largest genomic regions that lacked cacti 4 and 12 fragments were also assessed for enrichment in repetitive sequences. To identify repetitive elements, RepeatModeler v2.0.5 (Flynn et al. 2020) was run on the Galaxy Europe public server (The Galaxy Community et al. 2026), followed by RepeatMasker v4.1.5 (Smit et al. 2013). The proportion of each genomic region overlapping annotated repetitive sequences was then calculated. For comparison, repeat overlap was also quantified for all regions lacking cactus fragments, all cactus fragments, and the 0.1% largest cactus fragments.

Gene function within putatively introgressed regions (candidate Saguaro cacti 4 and 12) was evaluated using the R package SIGNET (Gouy et al. 2017), which identifies gene subnetworks enriched for high scores relative to a null model. Each of the 16,524 annotated genes was assigned an introgression score based on overlap with Saguaro-identified regions. Genes overlapping candidate cacti regions were assigned a score of 1, whereas genes lacking overlaps were scored as 0. These scores were used as input for SIGNET analyses. SIGNET was run separately using KEGG (Kanehisa and Goto 1999) and Reactome (Milacic et al. 2024) pathway databases. For each analysis, significance was assessed against a null distribution generated from 10,000 replicates, using a threshold of P < 0.01. A total of 1,546 Reactome pathways and 315 KEGG pathways were tested.

## Results and Discussion

### Sequencing and variant calling

Our contemporary genomic datasets comprised whole-genome sequencing data from 81 individuals, including all European species of hares (25 broom, 22 Italian, 6 mountain, 10 Iberian and 10 brown hares) and one outgroup (8 snowshoe hares) (Fig. 1A). Of these, 37 were newly sequenced for this work and the remainder were obtained from published datasets (Seixas et al. 2018; Giska et al. 2019; Souto et al. 2025) (Table S1). After quality filtering, the final SNP dataset including both high (54 genomes, 6 species; range: 6.43–24.74×) and low coverage (27 genomes, broom and Italian hares; range: 1.01–2.86×) samples contained 121,389,719 variants. Detailed sampling information, sequencing statistics and accession numbers are provided in Table S1. In addition, three ancient Italian hare mitochondrial genomes were obtained from Iron Age archaeological remains.

### Mitochondrial DNA replacement in the ancestor of broom and Italian hares

We were able to assemble the mitochondrial genomes of 76 contemporary individuals of five species and three Iron Age Italian hare samples. Their phylogeny was inferred based on 10,156 sites, resulting in the dated topology shown in Fig. S2, which is consistent with previous studies (Melo-Ferreira et al. 2012; Costa et al. 2024). Divergence times were also independently estimated within a fully Bayesian framework, restricted to the contemporary samples and thus relying solely on the deep node calibration (Supplementary Text S2; Fig. S3). Topology and relative node ages were highly consistent between the two approaches for the great majority of branches. We nonetheless favoured the least-squares approach for the main analysis, since combining a node calibration in the order of millions of years with tip dates only a few thousand years old within a single relaxed-clock Bayesian MCMC produced unstable rate estimates. This is a known limitation of Bayesian molecular dating, generally referred to as time-dependent rate bias (Ho et al. 2005; Ho et al. 2011). The least-squares approach, in contrast, natively accommodates such non-contemporaneous sampling. Combining both calibrations also allowed us to estimate the mitochondrial DNA substitution rate directly from the data. The inferred rate was higher than the nuclear-derived value assumed in a previous study (Seixas et al. 2018), consistent with the well-documented elevation of mitochondrial relative to nuclear substitution rates in mammals, which varies considerably across lineages (Allio et al. 2017).

These dates were transferred onto a broader mitochondrial phylogeny based on a cytochrome *b* fragment and including more samples (Costa et al. 2024), as well as several other species from the Palearctic and Nearctic close to the mountain hare and making up what we refer to as the “arctic clade” (Fig. 1B). Fig. 1B also shows the dated nuclear phylogeny of these species (adapted from Ferreira et al. 2021) for comparison. In brief, the “arctic” mitochondrial DNA clade includes recognized closely related arctic-boreal species (mountain hares, Alaskan hares and white-tailed jackrabbits) and haplotypes from several other species, reflecting many documented cases of mitochondrial DNA introgression of mountain hare origin (Melo-Ferreira et al. 2012; Seixas et al. 2018; Costa et al. 2024). Although the broom and Italian hares diverged from the arctic clade 2.39 Mya according to the nuclear phylogeny reported in Ferreira et al. (2021), they are included in this clade for mitochondrial DNA, with a maximum divergence of 0.89 Mya from all other members of the clade apart from the white-tailed jackrabbit (Fig. 1B).

This sharing of mitochondrial lineage suggests past massive mitochondrial DNA exchanges from mountain hares, which has been previously suggested by coalescent simulations (Melo-Ferreira et al. 2012). We will not discuss here the details of the results for species of the arctic clade other than the mountain hare, which given the biogeography and known evolutionary history of the “arctic” clade members is the likely donor. These other arctic-boreal species were here included to show that the discordance concerns the placement of broom-Italian hare haplotypes in the “arctic” clade, and thus that the direction of introgression was from mountain to broom-Italian hare ancestor rather than the reverse, which could have been equally inferred in the absence of native haplotypes if these other species were omitted. We will thus now concentrate only on the mountain hare, as the donor of mitochondrial DNA to broom-Italian hares.

In the mitochondrial DNA tree (Fig. 1B), the broom-Italian group appears paraphyletic relative to mountain hares. One basal lineage is specific to broom-Italian and diverged 0.89 Mya from a lineage that includes all mountain and some broom, as well as some Iberian and brown hare samples. The remaining brown and Iberian hare samples lie in branches of their own, external to the arctic clade. The position of these species-specific branches differs between the mitochondrial tree, where Iberian and brown are sisters, and the nuclear tree where they are not. However, previous work has shown that the mitochondrial DNA divergence between Iberian and brown hares is compatible with the nuclear DNA divergence, and this discordance can thus be attributed to lineage sorting (Melo-Ferreira et al. 2012). The other discordances between the two trees pointed above suggest complete fixation of mountain-related mitochondria in the ancestor of broom and Italian, at a date that must be at least 0.89 Mya (and 1.1 Mya, 95% HPD: 0.68-1.6 Mya, on the bayesian analysis), and more recent partial introgression that must have occurred in Iberia, since it affected the broom hare (but not the Italian hare), the Iberian hare and Iberian populations of the brown hare. Introgression in Iberia has been extensively documented elsewhere (Melo-Ferreira et al. 2005; Seixas et al. 2018) and we will concentrate here on the mitochondrial DNA replacement that affected the broom-Italian hare ancestor. The three ancient Italian hare samples nested within modern Italian hares (Fig. S21), and although based on a limited sample, this indicates that this mitochondrial lineage has persisted in this region since at least the Iron Age.

### Major pulse of genomic introgression into the ancestor of broom and Italian hares

Population structure based on autosomal sequences assessed within broom and Italian hares using NGSadmix (Skotte et al. 2013), suggested K=2 as the optimal number of genetic clusters (Fig. S4). While no clear spatial structure was detected in the broom hare, the Italian hare showed a pronounced north–south differentiation, consistent with restricted gene flow between regions (Fig. S4). This structure was considered in downstream analyses.

We compared 12 different models of the history of divergence and admixture between the mountain hare and the broom-Italian hare sisters (described in Fig. S5 and in Table S5) using BPP (Flouri et al. 2018), obtaining the likelihoods reported in Fig. S6. The model that best fitted the data according to AIC (mean log-likelihood = −1,594,375; 95% CI: −1,594,681 to −1,594,100; model XII in Fig. S5 and S4) involves pulses of introgression from the mountain hare into broom and Italian hares and their ancestor Fig. 1C. Under this model, divergence among the three species was estimated at 2.51 Mya (95% CI: 2.36–2.66 Mya), assuming a mutation rate of 2.8×10^-9^ substitutions/site/generation and a generation time of 2 years, which is consistent with previous phylogeny-based divergence estimates (Ferreira et al. 2021).

The pulse of introgression into the broom-Italian ancestor was dated ∼919 Kya (95% CI: 834 Kya–1.01 Mya). The dated mitochondrial tree independently placed the ancient mitochondrial DNA introgression at ∼0.89 Mya, remarkably close to this estimate and well within its 95% confident interval. The pulse of autosomal introgression was estimated to be substantial (φ = 24%; 95% CI: 19–29%). Souto et al. (2025) had reported only a modest contribution of the mountain hare to the genomes of the broom or Italian hares, in apparent contradiction with this result. This is because they estimated the mountain contribution independently in the broom and Italian hare, taking one or the other species as reference. They could thus only detect differential mountain hare introgression, i.e. that occurred after the divergence of broom and Italian hares or was differentially retained.

The split between broom and Italian hares was estimated at 64 Kya (95% CI: 57–72 Kya), also consistent with previous estimates (Ferreira et al. 2021; Souto et al. 2025), followed by minor pulses of mountain hare introgression ∼19 Kya (95% CI: 11–28 Kya) into the broom hare (φ = 1.08%) and ∼16 Kya (95% CI: 6–25 Kya) into the Italian hare (φ = 0.26%) (Table S4). The more pronounced recent introgression into the broom hare more than the Italian is consistent with the mitochondrial DNA results summarised above, since recent mitochondrial DNA introgression was detected in the broom but not in the Italian hare. This asymmetry is also consistent with the *D-statistics* estimates (D > 0, Z-score> 3; Table S6), and agrees with the low rates of introgression reported by Souto et al. (2025).

### Phylogenetic segmentation identifies introgressed genomic regions

According to the fitted demographic model, we expect about 24% of the genomes of broom and Italian hares to be of mountain origin. Given the estimated age of the admixture (about 5N generations given the estimated size of the ancestral population, Fig. 1C), we then expect about 76% of the broom-Italian hare genomes to have coalesced before the admixture (backwards in time). Given that the probability that the MRCA was of mountain hare origin is 0.24, we expect about 0.76*0.24 = 18% of the broom and Italian genomes to be fixed for mountain hare ancestry under the effect of random drift since the admixture. Note that some genomic regions could be fixed for mountain ancestry and still have an MRCA older than the introgression, but they should represent a small fraction of the expected 24% that did not coalesce before the introgression. Such genomic regions fixed for mountain ancestry should display a characteristic deviation from the species phylogeny, grouping together broom, Italian, and mountain hares.

The Saguaro method (Zamani et al. 2013) appears particularly suited to look for such genomic regions, since it is designed to partition the genome according to phylogenetic patterns, without the limitations of sliding window methods. After 13 cycles of Saguaro, the genome was partitioned into segments grouped in 13 “cacti”, i.e. distinct phylogenetic patterns (Fig. S7). One cactus was dominant (covering 41.54% of the genome) and matched the known species tree (Fig. 2A). We inspected the remaining cacti and identified cacti 4 and 12 showing a phylogeny matching extensive mountain introgression into broom-Italian hare (Fig. 2A). The segments belonging to cactus 4 spanned 17.3% of the genome, with median length 767 bp and containing a median of 37 SNPs per segment (Table S7; Fig. S8), while cactus 12 covered 0.5% of the genome, with segments of 119 bp median length and a median of 9 SNPs per segment (Table S8; Fig. S8). Together, cacti 4 and 12 segments covered ∼17.8% of the genome (Fig. S9), in good agreement with the 18% expectation based on the estimated parameters of the demographic model. The cactus 4 and 12 segments were spread across the genome, but some deserts of introgression are apparent visually (Fig. S9). The remaining cacti showed other possible phylogenies (not shown). Comparing the overlap of cacti 4 and 12 segments with low-mappability regions against a null distribution generated from 2,000 circular bootstrap replicates of the cactus fragments, showed that they were significantly depleted in low-mappability regions (Fig. S10), suggesting that their non-canonical phylogeny does not result from mapping artifacts. We also confirmed that the segments of these two cacti showed significantly lower Dxy between mountain hares and broom or Italian hares than expected under the null distribution generated using the circular bootstrap replicates (Fig. S11), which matches the expectations of introgression.

**Figure 2.**
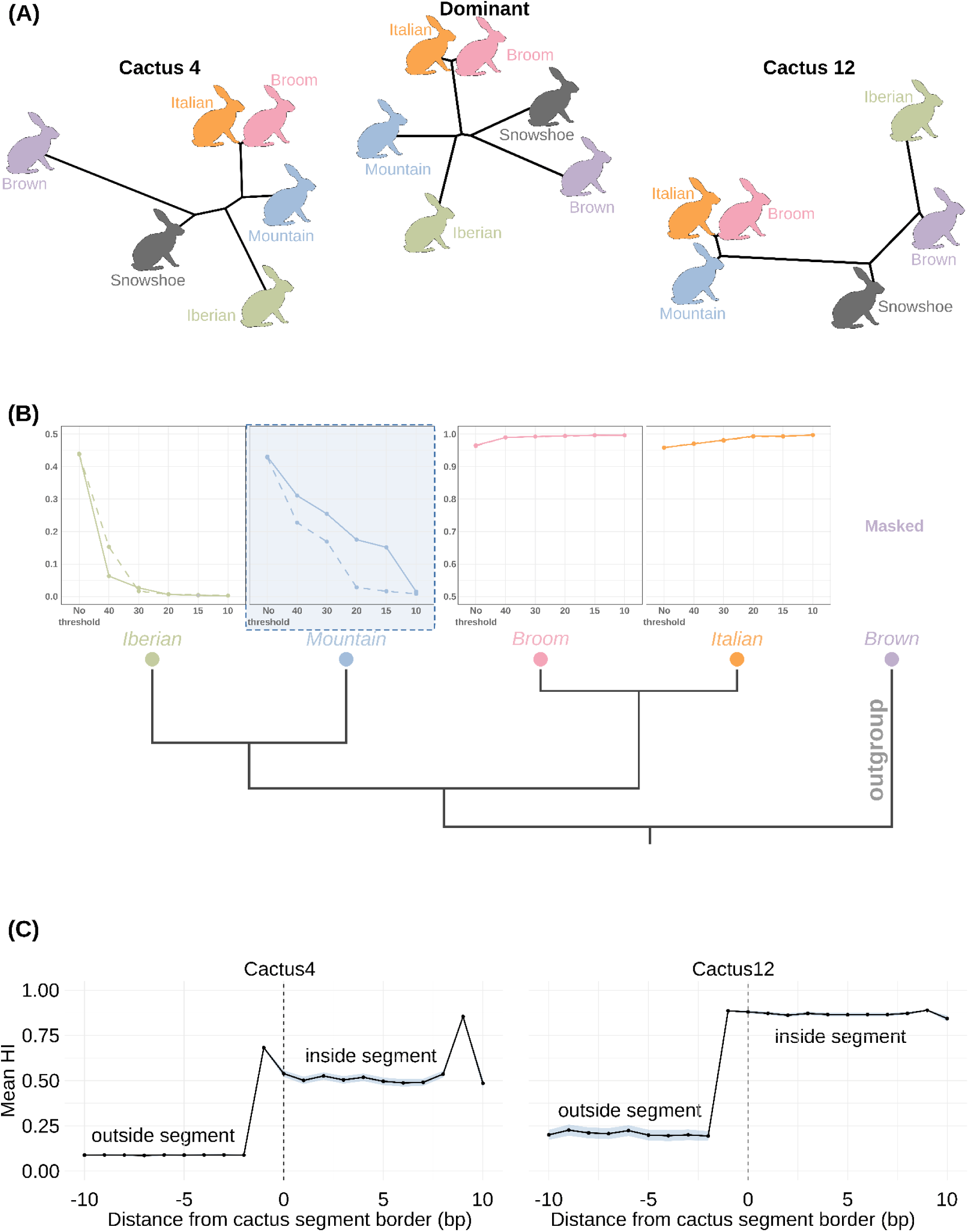
Characteristics of cacti 4 and 12. **(A)** phylogenetic patterns of cactus 4 (17.3% of the genome; left), the predominant cactus (41.5%; middle), and cactus 12 (0.51%; right). Both cactus 4 and cactus 12 are consistent with the inferred ancient introgression event, grouping broom, Italian, and mountain hares. **(B)** Mean hybrid index (HI) values estimated with *diem* in the different species, at different diagnosticity thresholds, for the whole autosomes (continuous curves) and excluding the cacti 4 and 12 segments (dashed curves). **C)** Variations of HI across the boundaries of cactus 4 and 12 segments in the mountain hare samples. The average HI of individual sites is plotted against their position relative to the segment boundary, with positive values inside the segment and negative values outside. Mean hybrid index (HI) across cactus boundaries for cactus 4 (left) and cactus 12 (right). Black lines represent the mean HI at each position, with grey shading indicating the 95% quantiles. Values are shown for the mountain hare. **Alt text:** Phylogenetic relationships in introgressed regions, *diem* polarization across different diagnostic indices, and Hybrid Index transitions across cactus barriers.

### Genome polarisation confirms highly introgressed regions

To explore the impact of mountain introgression more thoroughly into broom and Italian, we used genome polarisation (Baird et al. 2023; Setter et al. 2026). The method is intended to detect barriers to gene flow by polarising the alleles of biallelic sites so as to maximise overall linkage disequilibrium in a sample of genomes. It can also be used to quantify the amount of admixture across a barrier, if present. It is however not able to detect complete introgression from the genotypes representing only the donor (mountain hare in our case) and receiver (broom and Italian hares) species. We therefore added in the polarisation analysis a species that is sister to the donor, namely the Iberian hare, *L. granatensis* (Fig. 1B).

As expected, given the species tree (Fig. 1B), genome polarisation revealed a major genomic barrier between broom-Italian and mountain-Iberian hares. However, although hybrid indexes (HIs, at a diagnostic index, threshold of DI >= -20) were close to 1 for broom-Italian hares, and close to 0 for Iberian hares, mountain hare samples homogeneously displayed intermediate values (average HI = 0.283), suggesting that the barrier is partially broken between broom-Italian and mountain hares, as expected in an introgression scenario. However, this polarisation design cannot determine the direction of the exchanges, which could even be reciprocal. Here we interpret these results with the *a priori* that introgression occurred from mountain to broom-Italian hares, as explained above. Consequently, only the average values of HI in the mountain hare are relevant and represent the average effect of introgression present in the different samples of the true receivers (broom and Italian hares).

After removing fixed autapomorphies of the Iberian hare, which could be interpreted as admixture between mountain and broom-Italian hares, we tested different diagnosticity index (DI) thresholds (Fig. 2B). DI represents the likelihood of the observed genotypes at a given site across all samples, given the overall HIs of these samples. As there was very little variation of HI among conspecifics (not shown), we only show species averages. When the polarisation was done on the whole autosomes (continuous curves in Fig. 2B), broom and Italian hares displayed HIs close to 1 and Iberian hares close to 0, across the spectrum of DI thresholds. In contrast, mountain hares showed a gradual increase of HI with decreasing stringency of the DI threshold. When the polarisation was done excluding the fragments of cacti 4 and 12 (dashed curves in Fig. 2B), the profiles of broom-Italian and Iberian hares remained like those for the whole genome, while the mountain hare profile differed: HIs stayed close to 0 at high thresholds, increasing only at relatively low DI thresholds, indicating the strong contribution of the introgression cacti segments to the major signal of admixture. This was confirmed by showing that the average mountain HI in cacti 4 and 12 segments is considerably higher than the values obtained in circular bootstrap replicates of these cactus segments, especially for cactus 12 segments (Fig. S12).

To check the quality of the segmentation proposed by Saguaro, we inspected variations of HI across the borders of the cactus 4 and 12 segments. As shown in Fig. 2C, we observed a sharp transition of HI across the segment borders, with higher HI inside than outside. The values of HI varied depending on the DI threshold chosen, but the sharp transition at the borders remained whatever threshold was chosen (not shown). These results suggest that the Saguaro segmentation process correctly identified major transitions of ancestry along the genome, pinpointing segments of essentially mountain hare origin in broom and Italian hares.

The cacti we selected from the Saguaro analysis are then likely to represent genomic regions with high levels of introgression, since the phylogenetic signal they represent was consistently shared by all broom and Italian samples. In contrast, *diem* is expected to also detect incomplete introgression spread across samples and genomic regions. The sites of complete introgression are expected to exhibit the highest DI values after those of the perfectly diagnostic sites, especially if complete introgression is as prevalent as predicted by the Saguaro results (17.8%), or the theoretical expectation based on the demographic model (18%). The genome-wide distribution of DI values (Fig. S13) showed a pronounced peak near 0 (corresponding to perfectly diagnostic sites), alongside a secondary peak around DI ≈ -15, which would correspond to completely introgressed sites. In the mountain hare HI profile for whole-genome polarisation (of Fig. 2B), a plateau of HI between 15 and 18% was found at the highest DI thresholds (LL -15 to -20). This likely corresponds to the contribution of fixed introgression, pointing to a genomic fraction in remarkable agreement with the two other estimates. In line with this, we found that cacti 4 and 12 segments have a much higher density of high-DI sites than their circular bootstrap replicates (Fig. S14). At less stringent DI thresholds (LL -30 to - 40) one should recover effects of non-fixed introgression, and we find HI values of 25-30% when polarising the whole genome, in good agreement with the prediction of the demographic model (24%, 95% CI: 19-29%).

Although neither *diem* nor Saguaro is a real ancestry deconvolution method based on a model of coalescent with recombination, their results remarkably agree, despite using completely different aspects of the data. It is therefore likely that we were able to find true segments of introgression, but only a model-based method could allow evaluating this hypothesis. Even so, we have demonstrated here the successful use of two non-canonical methods in the context of the detection of introgression. They are both model-free and use simple aspects of the data, genetic distance for Saguaro and linkage disequilibrium for *diem*. The former uses the physical position of sites along the genome, while the second does not. Yet they remarkably agree, as well as with the results of a more complex and model-based method based on the coalescent, but on a small subsample of the genome. A combination of the former two methods could be an interesting perspective, *i.e.,* segmenting the genome *à la* Saguaro based on the site information of *diem*. More work would also be needed to determine the possibilities and conditions of application of *diem* when including a sister taxon to the donor in the analysis.

Despite this strong concordance between Saguaro and *diem*, both analyses were performed on a subset of individuals. To assess the extent to which the inferred cactus 4 and cactus 12 segments were fixed across all sampled individuals, we generated SNP heatmaps for two datasets: one including only the high-coverage samples and a second including all available broom and Italian hare samples. For each SNP, the majority allele in the broom hare was defined as the reference allele, and genotypes were classified as homozygous reference, heterozygous, or homozygous alternative. We then calculated the proportion of non-missing sites that were homozygous for the reference allele in broom and Italian hares in each dataset. This proportion was highly consistent between datasets: 97.2% in the high-coverage dataset and 98.1% when all samples were included (Tables S9 and S10). These results suggest that the inferred cactus segments are nearly fixed across the sampled broom and Italian hare populations, with no evidence that the inclusion of additional, lower-coverage individuals altered the inferred pattern.

### Repeatability of introgressed genes across biological systems

We then confronted our results to those of Seixas et al. (2018), who inferred introgression of the mitochondrial and nuclear genes from the mountain into the Iberian hare, and suggested candidate genes for adaptive introgression. That work detected nuclear introgression based on Dxy and pinpointed genes overlapping genomic windows with higher frequency introgression than expected under a null demographic model, a pattern expected in case of adaptive introgression. It also pinpointed nuclear genes with mitochondrial functions whose geographic gradient of introgression paralleled that observed for mitochondrial DNA introgression, a pattern expected in case of nucleo-cytoplasmic coadaptation driving co-introgression. We will here call all these genes “candidate genes”. Of the 86 such genes with unambiguous annotations that could be matched between the annotation systems used in Seixas et al. (2018) and the present study, 77 overlapped cactus 4 or cactus 12 segments. Among the overlapping genes, 59 had coding sequences (CDS) overlapping these cactus segments. We found, however, that these seemingly high proportions are not unexpected when considering genome coverage: the proportion of candidate gene coverage relative to total gene coverage within cactus 4 and 12 regions did not differ from expectations under the circular bootstrap null distribution (coverage ratio = 0.01; p = 0.48; Fig. S15). This reflects the fact that cactus 4 is enriched for both genes and coding sequences (CDS), as also revealed by circular bootstrap sampling (Fig. S16; cactus 12 segments were not included here since they represent a very small genomic fraction). Therefore, the high proportion of candidate genes (*sensu* Seixas et al. 2018) overlapping cactus 4 and 12 segments may be due to these segments being enriched in genes, and no link can be currently established between the nature of these genes and their level of introgression on this basis.

Despite this, eight genes with annotated mitochondrial functions were identified within introgressed coding regions: MRPL13, TUFM, PITRM1, MCCC1, HCLS1, CLYBL, HEBP1, and RPL34. However, pathway-level enrichment analyses did not identify any significantly overrepresented mitochondria-related pathways among the KEGG or Reactome subnetworks detected (Fig. S17). This may reflect the functional diversity of mitochondrial genes, which participate in processes ranging from mitochondrial translation and respiratory chain assembly to intermediary metabolism, preventing them from forming a single topologically coherent subnetwork detectable by SIGNET. Alternatively, these genes may simply represent too small a fraction of the ∼531 MitoCarta-annotated genes overlapping introgressed regions to achieve genome-wide significance. Notably, eight of the thirteen mitochondrial-function genes highlighted by Seixas et al. (2018) (Table S9) as exhibiting introgression patterns paralleling the geographic distribution of mitochondrial introgression from the mountain hare into the Iberian hare were also identified in our analyses. Although these genes are not structural components of the oxidative phosphorylation complexes, they perform key mitochondrial functions, including translation, respiratory chain assembly, and metabolism (see Seixas et al. 2018 and reference therein). Their repeated association with introgressed regions across independent systems is consistent with the hypothesis that mitonuclear interactions contribute to the preferential retention of introgressed variation. The comparison between our results and those of Seixas et al. (2018) has limitations, since it is based on the intersection of gene annotations rather than genomic coordinates. This is because Seixas et al. detected introgression using a window-based approach, while we did not. Using similar methods for the two systems would allow a more rigorous comparison and should be the object of further investigation.

### Genomic and functional characteristics of highly and poorly introgressed regions

We additionally inspected some characteristics of the genes overlapping the highly introgressed segments of cacti 4 and 12. We identified 531 genes with overlapping CDS and with potential mitochondrial function according to Mitocarta 3.0 (Rath et al. 2021). We reasoned that if strong selection had favoured the introgression of some genes, they should tend to lie in relatively long introgression segments that quickly became fixed by selection before being broken by recombination. The largest cactus 4 or 12 fragments were thus examined for the presence of genes of interest. For cactus 4, the 0.1% largest fragments comprised 31 regions ranging from 29 to 100 kb. These regions overlapped 24 genes, of which only six had annotated gene IDs (ACAD11, ACKR4, TDRD1, TNIK, TTC21B, and ZNF654). Among these, only ACAD11 has a reported mitochondrial function being involved in mitochondrial fatty acid metabolism and is thus another candidate inviting us to consider the hypothesis that mitonuclear genes may contribute to the maintenance of introgressed mitochondrial genomes. Of the remaining 18 genes, 14 were successfully annotated by homology using BLAST. These genes are primarily associated with transcriptional regulation, olfactory receptors, immune function, and spermatogenesis, functional categories that have previously been implicated in adaptive introgression across mammals, including hares (Seixas et al. 2018; Silvert et al. 2019; Banker et al. 2022). Functional descriptions of all annotated genes are provided in Table S11. These functional categories represent plausible targets of adaptive introgression, either by facilitating compatibility with the foreign genomic background or by contributing to adaptation to novel environmental conditions, as already argued in Seixas et al. (2018), who found an enrichment of these categories among highly introgressed genes in the Iberian hare. For cactus 12, the 0.1% largest fragments comprised six regions ranging from 11 to 20 kb. A single gene overlapped these fragments, and although no gene ID could be assigned, it was successfully annotated by homology using BLAST.

We then also examined genomic regions devoid of introgressed cactus 4 or 12 segments (Fig. S9). The 0.1% largest segments comprised 222 regions between 55 and 447 kb. Details of these regions are provided in Table S12. Gene Ontology (GO) enrichment analysis of the genes overlapping these regions identified a single significantly enriched functional category, the NF-kappa B signalling pathway, which regulates the survival, activation and differentiation of innate immune cells and inflammatory T cells (Liu et al. 2017). Across chromosomes, chromosome 21 contained the lowest proportion of cactus regions, whereas chromosome 3 showed the greatest extent of inferred introgression. These fragments showed a high proportion of repetitive DNA, with approximately 83% of their sequence overlapping repetitive elements identified by RepeatMasker. This value was higher than the corresponding overlap for all regions lacking cactus 4 and 12 fragments (41%), or for cactus 4 and 12 regions (38%).

Overall, the observation of an enrichment in coding sequences in highly introgressed regions and in repeated sequences in less introgressed regions may reflect correlations of these quantities with local recombination rates. In the presence of incompatibilities between the genomes of the two species, introgression is facilitated when recombination breaks linkage to such incompatibilities (Martin and Jiggins 2017). It has been generally observed across various taxa that high recombination regions generally tend to be more gene-dense and low recombination regions more repeat-rich (Tiley and Burleigh 2015; Kent et al. 2017; Stapley et al. 2017). Provided such correlation also holds in hares, for which a direct recombination map is not yet available, the recombination landscape could account for variations of introgression along the autosomes. However, we cannot exclude a confounding effect, since repeats may negatively affect the quality of SNP calling, and thus the ability to reliably detect introgression.

One important prospect for future research concerns the sex chromosomes, which were not included in the present study. Based on common observations, we expect to see less introgression on the X than on the autosomes, because of the well-documented large X effect (Presgraves 2018). Verifying this would be a confirmation that genetic incompatibilities exist between these species, which is highly expected. The only indication we have of this with the autosomes is the very indirect evidence of a correlation between recombination and introgression.

## Conclusion

In this work, we confirmed that a complete mitochondrial replacement occurred in the common ancestor of two sister hare species (broom and Italian) and was accompanied by nuclear introgression affecting a substantial fraction of the genomes, of the order of 18%, distributed in numerous spread short fragments spread and reaching fixation. This estimate emerged consistently from three independent lines of evidence, coalescent demographic modelling, phylogenetic segmentation, and genome polarisation. Despite relying on markedly different aspects of the data, the convergence of results lends confidence both to the extent of introgression inferred and to the combined approach itself, as a general strategy for detecting extensive or even fixed introgression in the absence of non-introgressed reference populations. Such massive genomic exchanges can have a significant impact on other types of inference: in our data, this single admixture pulse alone is enough to place the broom-Italian ancestor artefactually within the arctic clade in the mitochondrial tree, despite the two lineages having diverged 2.39 Mya at the nuclear level, and at regions along the genome. Since mitochondrial introgression from the mountain hare is widespread across several hares species, a similar assessment of nuclear introgression, e.g. using the combined approach applied here, would be needed to establish how general this pattern is, and might invite a revision of the dated phylogeny of the genus.

## Acknowledgments

We thank Istituto Superiore per la Protezione e la Ricerca Ambientale (ISPRA) for providing the Italian hare samples utilized in this work. This work was supported by project HybridChange (doi: 10.54499/PTDC/BIA-EVL/1307/2020), funded by Fundação para a Ciência e a Tecnologia (FCT). J.C. was supported by an FCT doctoral scholarship (UI/BD/154450/2022). Computational resources were provided through the FCT Advanced Computing Projects 2025.00006.CPCA and 2025.09424.CPCA. The work was further supported by the BIOPOLIS Education, Research and Outreach Program (project LEPINTRO), the European Union’s Horizon 2020 research and innovation programme under grant agreement No 857251, and by the Regional NORTE 2030 Program via the project NORTE2030-FEDER-02130700.

## Author Contributions

J.M.-F., J.P.M., P.B., and J.C. conceived and designed the study. J.M.-F., J.P.M., and P.B. supervised the work. J.C. and J.P.M. conducted the analyses. P.C.A and J.Q. led sampling of contemporary specimens. L.F. performed modern DNA laboratory work. A.T. led sampling of ancient specimens; S.G.M. performed the ancient DNA laboratory work; E.S. analysed ancient DNA data. G.L. and J.A. supervised the ancient DNA component of the study. J.C., J.P.M., P.B., and J.M.-F. wrote the manuscript. All authors reviewed and approved the final manuscript.

## Data availability

All code used in data analyses and visualization is available in GitHub at: https://github.com/evochange/Ancient-Shared_Introgression_Hares. The sequencing dataset supporting the conclusions of this article is available in the National Centre for Biotechnology Information (NCBI) under the accession numbers PRJNA1518870, PRJNA399194, PRJNA562432, PRJNA561428, PRJNA561582, PRJNA564335 and PRJNA420081.

