## Supplementary Material for "Extensive nuclear introgression accompanied ancient mitochondrial capture in the common ancestor of two sister hare species"

### Supplementary Text

*Text S1 – Ancient DNA sampling, laboratory procedures and bioinformatic processing*

Three Italian hare bone samples used for ancient DNA analysis originate from Cavità 254, a stone quarry at Orvieto (Italy) that was rapidly infilled during the late Iron Age (Etruscan period; Trentacoste 2021) (Fig. S1). The hare bones themselves were not directly radiocarbon dated; the chronology of the deposit is instead based on direct AMS dating of an associated chicken bone from the same stratigraphic context (775–541 BC, 95% confidence; Best et al. 2022), consistent with the broader chronology of the bone-rich deposit, which was well sealed (George et al. 2017). Seven hare bone samples from this context were initially processed for ancient DNA. Extracted DNA was shotgun-sequenced at low coverage to assess preservation; the proportion of endogenous hare DNA ranged from 4.8% to 14.5% across the seven libraries. Based on these results, three libraries, hereafter Lcor Iron 125, Lcor Iron 127, and Lcor Iron 129, were selected for downstream mitochondrial target enrichment.

All laboratory procedures were carried out in a dedicated ancient DNA facility at the Palaeogenomics & Bio-Archaeology Research Network, University of Oxford, following standard contamination-control measures (positive air pressure, UV irradiation, bleach decontamination, disposable protective clothing). DNA was extracted following a modified version of the protocols of Dabney et al. (2013) and Damgaard et al. (2015), with extraction negative controls included throughout. Double-stranded sequencing libraries were prepared following Carøe et al. (2018), with the number of indexing PCR cycles determined by qPCR; negative controls were again included during library preparation. Libraries were first screened by low-coverage paired-end sequencing (2×150 bp; Illumina HiSeq 4000, Novogene, Cambridge, UK) to estimate endogenous DNA content and guide pooling. The three selected libraries were then enriched for the mitochondrial genome using custom in-solution hybridization probes (MyBaits, Arbor Biosciences) targeting the brown hare mitogenome, and the captured libraries were sequenced on an Illumina HiSeq X platform (2×150 bp; Macrogen, Seoul, South Korea).

Sequencing reads were processed with AdapterRemoval v2.3.2 (Schubert et al. 2016), removing adapter sequences, quality-filtering reads (--trimns, --trimqualities), and merging overlapping read pairs (-collapse, --preserve5p). Merged, unmerged, and singleton reads were mapped independently with BWA aln v0.7.17 (Li and Durbin 2009; seeding disabled, -l 1024; -n 0.01; -o 2), a parameter combination shown to reduce reference bias in ancient DNA mapping at increased computational cost (Oliva et al. 2021). Reads were mapped against a hare pseudogenome (Marques et al. 2020) in which the mitochondrial sequence was replaced with an Italian hare mitogenome (GenBank accession KJ397606), using the CircularMapper module of EAGER (Peltzer et al. 2016) to account for the circularity of the mitochondrial reference, with a 500-bp extension across the artificial breakpoint. PCR duplicates were removed with Picard MarkDuplicates v2.18.23, and consensus mitochondrial sequences were generated with htsbox pileup (r345; minimum read length 30 bp, mapping and base quality ≥30, minimum coverage 5×, majority-allele consensus). Mitochondrial genome coverage was calculated with GATK v3.8 (McKenna et al. 2010); minimum mapping and base quality 30). The three consensus mitochondrial genomes achieved mean coverages of 53.4×, 63.3×, and 141.7×, with 3.0%, 2.4%, and 2.5% missing data, respectively.

*Text S2 – Confirmatory dating analysis using contemporary samples only*

To confirm the divergence times inferred from the combined ancient-modern mitochondrial dataset (main text), we independently estimated the same divergence times using a fully Bayesian approach restricted to the 76 contemporary samples. This analysis avoids the need to reconcile calibrations spanning very different temporal scales, the deep snowshoe–brown hare node (Mya) and the ancient tip dates (years), within a single relaxed-clock MCMC, which we found to produce unstable rate estimates when both types of calibration were combined (see main text).

The 13 mitochondrial protein-coding genes and two rRNA genes were concatenated and aligned using MAFFT v7.505 (Katoh and Standley 2013). The concatenated alignment was analyzed in BEAST v2.7.8 (Bouckaert et al. 2019) using two partitions (protein-coding and rRNA genes), with unlinked substitution and clock models and a linked tree model. Both partitions were modeled under GTR with a gamma category count set to 4.0 and a relaxed lognormal clock. A calibrated Yule prior was applied, with the snowshoe–brown hare MRCA constrained to 3.99 Mya (95%

HPD: 3.59–4.36 Mya; Ferreira et al. 2021). Markov chain Monte Carlo (MCMC) analyses were run for 200 million generations with three independent replicates. Convergence and mixing were assessed in Tracer v1.7.2 (Rambaut et al. 2018). Runs were combined using LogCombiner v2.7.8 (Drummond and Rambaut 2007), and posterior trees were summarized into a maximum clade credibility tree using TreeAnnotator v2.7.8 (Drummond and Rambaut 2007). The resulting tree (Fig. S3) was visualized in FigTree v1.4.4 (Rambaut 2018).

### Supplementary Tables

*Table S1 – Sample Info*

| *ID* | *Species* | *Population* | *Country* | *Reference* | *BioSample Acession* | *Mean Depth* | *Mean GQ* |
|---|---|---|---|---|---|---|---|
| *Lgra1* | *granatensis* | Alcoutim | Portugal | Seixas et al. 2018 | SAMN07526963 | 20,9174 | 87.7433 |
| *Lgra2* | *granatensis* | Peñaflor | Spain | Seixas et al. 2018 | SAMN07526964 | 20,0237 | 86.2217 |
| *Lgra3* | *granatensis* | Pancas | Portugal | Seixas et al. 2018 | SAMN07526965 | 17,1856 | 82.0126 |
| *Lgra4* | *granatensis* | Idanha | Portugal | Seixas et al. 2018 | SAMN07526966 | 22,1227 | 91.0585 |
| *Lgra5* | *granatensis* | Miguelturra | Spain | Seixas et al. 2018 | SAMN07526967 | 22,451 | 92.0394 |
| *Lgra6* | *granatensis* | Valpaços | Portugal | Seixas et al. 2018 | SAMN07526968 | 22,668 | 91.434 |
| *Lgra7* | *granatensis* | Algete | Spain | Seixas et al. 2018 | SAMN07526969 | 21,8257 | 90.4382 |
| *Lgra8* | *granatensis* | Valência | Spain | Seixas et al. 2018 | SAMN07526970 | 17,7685 | 82.9866 |
| *Lgra9* | *granatensis* | Monte Allá Detrás-Sauguillo. Torrubia de Soria | Spain | Seixas et al. 2018 | SAMN07526971 | 16,8502 | 82.4207 |
| *Lgra10* | *granatensis* | Fontellas | Spain | Seixas et al. 2018 | SAMN07526972 | 23,6069 | 92.8816 |
| *Lcas1* | *castroviejoi* | Cospedal | Spain | Souto et al. 2025 | SAMN12640761 | 16,418 | 99.2963 |
| *Lcas2* | *castroviejoi* | Alto Sil - León | Spain | Souto et al. 2025 | SAMN12640762 | 10,5262 | 94.0878 |
| *Lcas3* | *castroviejoi* | Alto Sil | Spain | Souto et al. 2025 | SAMN12640763 | 7,63282 | 89.6312 |
| *Lcas4* | *castroviejoi* | León | Spain | Souto et al. 2025 | SAMN12640764 | 6,83025 | 88.683 |
| *Lcas5* | *castroviejoi* | León | Spain | Souto et al. 2025 | SAMN12640765 | 8,20967 | 91.1067 |
| *Lcas6* | *castroviejoi* | Villablino (Sierra de Omaña) | Spain | This work | SAMN62699280 | 12,1988 | 96.6257 |
| *Lcas7* | *castroviejoi* | Villablino (Sierra de Omaña) | Spain | This work | SAMN62699281 | 11,8253 | 96.0697 |
| *Lcas8* | *castroviejoi* | Salce | Spain | This work | SAMN62699282 | 9,4686 | 93.165 |
| *Lcas9* | *castroviejoi* | Cantabria | Spain | This work | SAMN62699283 | 11,2428 | 95.6673 |
| *Lcas19* | *castroviejoi* | Alto Sil | Spain | This work | SAMN62699284 | 9,62212 | 93.7732 |
| *Lcas20* | *castroviejoi* | Alto Sil | Spain | This work | SAMN62699285 | 1,70301 | 64.2412 |
| *Lcas21* | *castroviejoi* | Alto Sil | Spain | This work | SAMN62699286 | 1,97961 | 70.1554 |
| *Lcas22* | *castroviejoi* | Alto Sil | Spain | This work | SAMN62699287 | 1,3344 | 59.6329 |
| *Lcas10* | *castroviejoi* | El Acebo | Spain | This work | SAMN62699288 | 1,59715 | 64.327 |
| *Lcas11* | *castroviejoi* | Asturias | Spain | This work | SAMN62699289 | 1,37272 | 60.3493 |
| *Lcas12* | *castroviejoi* | Argovejo | Spain | This work | SAMN62699290 | 1,57036 | 63.6952 |
| *Lcas13* | *castroviejoi* | Cantabria | Spain | This work | SAMN62699291 | 2,22352 | 73.53 |
| *Lcas14* | *castroviejoi* | Riano | Spain | This work | SAMN62699292 | 2,83308 | 78.6552 |
| *Lcas15* | *castroviejoi* | León | Spain | This work | SAMN62699293 | 2,86301 | 79.1195 |
| *Lcas16* | *castroviejoi* | San Emiliano | Spain | This work | SAMN62699294 | 2,84464 | 78.8819 |
| *Lcas17* | *castroviejoi* | Cangas de Narcea | Spain | This work | SAMN62699295 | 1,47795 | 62.7755 |
| *Lcas18* | *castroviejoi* | Alto Sil - León | Spain | This work | SAMN62699296 | 1,76945 | 67.8667 |

(continue…)

| *ID* | *Species* | *Population* | *Country* | *Reference* | *BioSample Acession* | *Mean Depth* | *Mean GQ* |
|---|---|---|---|---|---|---|---|
| *Leur1* | *europaeus* | Cantabria | Spain | Seixas et al. 2017 | SAMN12618114 | 20,6963 | 78.0876 |
| *Leur2* | *europaeus* | Villareal del Canal | Spain | Seixas et al. 2017 | SAMN12618115 | 22,7387 | 83.2835 |
| *Leur3* | *europaeus* | Villarcayo | Spain | Seixas et al. 2017 | SAMN12618116 | 21,0113 | 77.3475 |
| *Leur4* | *europaeus* | Álava | Spain | Seixas et al. 2017 | SAMN12618117 | 19,4759 | 75.1883 |
| *Leur5* | *europaeus* | Alsasua | Spain | Seixas et al. 2017 | SAMN12618118 | 21,7109 | 75.5745 |
| *Leur6* | *europaeus* | Pyrenees | France | Seixas et al. 2017 | SAMN12618119 | 21,5852 | 80.8384 |
| *Leur7* | *europaeus* | Oblast Kiev | Ukraine | Seixas et al. 2017 | SAMN12618120 | 17,1739 | 70.7522 |
| *Leur8* | *europaeus* | Germany | Germany | Seixas et al. 2017 | SAMN12618121 | 14,0654 | 58.2338 |
| *Leur9* | *europaeus* | Vienna | Austria | Seixas et al. 2017 | SAMN12618122 | 15,6997 | 64.549 |
| *Leur10* | *europaeus* | Clermont-Ferrand | France | Seixas et al. 2017 | SAMN12618123 | 19,7684 | 76.0979 |
| *Ltim1* | *timidus* | Borris-in-Ossory | Ireland | Marques et al. 2019 | SAMN12621015 | 22,5348 | 88.6838 |
| *Ltim2* | *timidus* | Captivity | Finland? | Seixas et al. 2018 | SAMN07526960 | 19,549 | 84.1836 |
| *Ltim3* | *timidus* | Calfreisen. Egga | Switzerland | Seixas et al. 2018 | SAMN07526961 | 21,1592 | 88.8934 |
| *Ltim4* | *timidus* | Nancy-sur-Cluses | France | Seixas et al. 2018 | SAMN07526962 | 24,7394 | 94.8513 |
| *Ltim5* | *timidus* | Signabøur | Faroe Islands | Giska et al. 2019 | SAMN12710256 | 7,52242 | 58.6284 |
| *Ltim6* | *timidus* | Amurskaya territory. Norsky reserve | Russia | This work | SAMN62699317 | 21,5478 | 92.2096 |
| *Lcor1* | *corsicanus* | Córsega | France | Souto et al. 2025 | SAMN12640766 | 20,6884 | 98.8292 |
| *Lcor2* | *corsicanus* | Córsega | France | Souto et al. 2025 | SAMN12640767 | 10,2593 | 88.7674 |
| *Lcor3* | *corsicanus* | Córsega | France | Souto et al. 2025 | SAMN12640768 | 10,6712 | 88.7039 |
| *Lcor4* | *corsicanus* | Córsega | France | Souto et al. 2025 | SAMN12640769 | 9,27796 | 87.7097 |
| *Lcor5* | *corsicanus* | Córsega | France | Souto et al. 2025 | SAMN12640770 | 10,5746 | 88.9287 |
| *Lcor6* | *corsicanus* | National Park Pollino | Italy | This work | SAMN62699297 | 13,1844 | 94.6451 |
| *Lcor7* | *corsicanus* | Tolfa (Roma) | Italy | This work | SAMN62699298 | 9,57178 | 89.0579 |
| *Lcor8* | *corsicanus* | Lazio | Italy | This work | SAMN62699299 | 16,3865 | 97.9305 |
| *Lcor9* | *corsicanus* | National Park Circeo | Italy | This work | SAMN62699300 | 9,05133 | 88.4773 |
| *Lcor10* | *corsicanus* | Lucanian Dolomites | Italy | This work | SAMN62699301 | 10,0944 | 90.5972 |
| *Lcor11* | *corsicanus* | National Park Pollino | Italy | This work | SAMN62699302 | 1,45992 | 58.3496 |
| *Lcor12* | *corsicanus* | National Park Pollino | Italy | This work | SAMN62699303 | 1,3675 | 56.5137 |
| *Lcor13* | *corsicanus* | National Park Pollino | Italy | This work | SAMN62699304 | 1,23494 | 53.7777 |
| *Lcor14* | *corsicanus* | National Park Pollino | Italy | This work | SAMN62699305 | 1,06008 | 49.2677 |
| *Lcor15* | *corsicanus* | National Park Pollino | Italy | This work | SAMN62699306 | 1,68851 | 62.3756 |
| *Lcor16* | *corsicanus* | National Park Pollino | Italy | This work | SAMN62699307 | 1,50892 | 59.1084 |
| *Lcor17* | *corsicanus* | National Park Pollino | Italy | This work | SAMN62699308 | 1,21668 | 53.1507 |

(continue…)

| ID | Species | Population | Country | Reference | BioSample Acession | Mean Depth | Mean GQ |
|---|---|---|---|---|---|---|---|
| *Lcor18* | *corsicanus* | Terni | Italy | This work | SAMN62699309 | 1,01259 | 47.8953 |
| *Lcor19* | *corsicanus* | Lazio | Italy | This work | SAMN62699310 | 1,20363 | 52.8329 |
| *Lcor20* | *corsicanus* | Lazio | Italy | This work | SAMN62699311 | 1,07464 | 49.4492 |
| *Lcor21* | *corsicanus* | Tolfa (Roma) | Italy | This work | SAMN62699312 | 1,38281 | 56.6372 |
| *Lcor22* | *corsicanus* | Tolfa (Roma) | Italy | This work | SAMN62699313 | 1,01687 | 48.0972 |
| *Lcor23* | *corsicanus* | Tolfa (Roma) | Italy | This work | SAMN62699314 | 1,51479 | 59.4714 |
| *Lcor24* | *corsicanus* | Lucanian Dolomites | Italy | This work | SAMN62699315 | 1,51191 | 59.1084 |
| *Lcor25* | *corsicanus* | Manciano (Grosseto) | Italy | This work | SAMN62699316 | 1,26823 | 54.7462 |
| *Lame1* | *americanus* | Near Lake Inez. Missoula | USA | Seixas et al. 2018 | SAMN07526959 | 22,8176 | 84.5529 |
| *Lame2* | *americanus* | Boreal | USA | Jones et al. 2018 | SAMN13999397 | 14,9331 | 69.4581 |
| *Lame3* | *americanus* | Pacific Northwest | Canada | Jones et al. 2018 | SAMN08146526 | 9,47961 | 55.3295 |
| *Lame4* | *americanus* | Rocky Mountains | USA | Jones et al. 2018 | SAMN08146529 | 6,43374 | 41.8676 |
| *Lame5* | *americanus* | Boreal | USA | Jones et al. 2018 | SAMN08146528 | 6,68557 | 46.1132 |
| *Lame6* | *americanus* | Pacific Northwest | USA | Jones et al. 2018 | SAMN08146466 | 13,5247 | 65.33 |
| *Lame7* | *americanus* | Pacific Northwest | USA | Jones et al. 2018 | SAMN08146494 | 23,7582 | 86.7425 |
| *Lame8* | *americanus* | Boreal | USA | Jones et al. 2018 | SAMN19037327 | 11,111 | 57.9684 |
| *Lcor Iron 125* | *corsicanus* | Ancient | Italy | This work | NA | mtDNA only | mtDNA only |
| *Lcor Iron 127* | *corsicanus* | Ancient | Italy | This work | NA | mtDNA only | mtDNA only |
| *Lcor Iron 129* | *corsicanus* | Ancient | Italy | This work | NA | mtDNA only | mtDNA only |

*Table S2 – Mitogenome Assembly*

| ***Sample*** | ***Mitogenome Assembly tool*** | ***Subset coverage*** | ***Complete Mitochondria Assembled*** | ***NCBI Acession Number*** |
|---|---|---|---|---|
| *Lcas1* | MitoCatch | 0.1 | YES | PZ900416 |
| *Lcas2* | MitoCatch | 0.3 | YES | PZ900420 |
| *Lcas3* | NOVOPlasty (Dierckxsens et al. 2016) | - | YES | PZ900421 |
| *Lcas4* | MitoCatch | 0.1 | YES | PZ900422 |
| *Lcas5* | MitoCatch | 0.1 | YES | PZ900423 |
| *Lcas6* | MitoCatch | 0.1 | YES | PZ900424 |
| *Lcas7* | MitoCatch | 0.3 | YES | PZ900425 |
| *Lcas8* | MitoCatch | 0.3 | YES | PZ900426 |
| *Lcas9* | MitoCatch | 0.3 | YES | PZ900427 |
| *Lcas10* | MitoCatch | 0.1 | YES | PZ900406 |
| *Lcas11* | MitoCatch | 0.1 | YES | PZ900407 |
| *Lcas12* | MitoCatch | 0.1 | YES | PZ900408 |
| *Lcas13* | MitoCatch | 0.3 | YES | PZ900409 |
| *Lcas14* | NOVOPlasty (Dierckxsens et al. 2016) | - | YES | PZ900410 |
| *Lcas15* | MitoCatch | 0.1 | YES | PZ900411 |
| *Lcas16* | MitoCatch | 0.1 | YES | PZ900412 |
| *Lcas17* | MitoCatch | 0.1 | YES | PZ900413 |
| *Lcas18* | MitoCatch | 0.1 | YES | PZ900414 |
| *Lcas19* | MitoCatch | 0.1 | YES | PZ900415 |
| *Lcas20* | MitoCatch | 0.1 | YES | PZ900417 |
| *Lcas21* | MitoCatch | 0.1 | YES | PZ900418 |
| *Lcas22* | MitoCatch | 0.3 | YES | PZ900419 |
| *Lcor1* | MitoCatch | 0.1 | YES | PZ900438 |
| *Lcor2* | MitoCatch | 0.1 | YES | PZ900444 |
| *Lcor3* | MitoCatch | 0.1 | YES | PZ900445 |
| *Lcor4* | MitoCatch | 0.1 | YES | PZ900446 |
| *Lcor5* | MitoCatch | 0.1 | YES | PZ900447 |
| *Lcor6* | MitoCatch | 0.1 | YES | PZ900448 |
| *Lcor7* | NOVOPlasty (Dierckxsens et al. 2016) | - | NO | - |
| *Lcor8* | MitoCatch | 0.1 | YES | PZ900449 |
| *Lcor9* | MitoCatch | 0.1 | YES | PZ900450 |
| *Lcor10* | MitoCatch | 0.1 | YES | PZ900428 |
| *Lcor11* | MitoCatch | 0.1 | YES | PZ900429 |
| *Lcor12* | MitoCatch | 0.1 | YES | PZ900430 |
| *Lcor13* | MitoCatch | 0.1 | YES | PZ900431 |
| *Lcor14* | MitoCatch | 0.1 | YES | PZ900432 |
| *Lcor15* | MitoCatch | 0.1 | YES | PZ900433 |
| *Lcor16* | MitoCatch | 0.1 | YES | PZ900434 |
| *Lcor17* | MitoCatch | 0.1 | YES | PZ900435 |
| *Lcor18* | MitoCatch | 0.1 | YES | PZ900436 |
| *Lcor19* | MitoCatch | 0.1 | YES | PZ900437 |
| *Lcor20* | MitoCatch | 0.1 | YES | PZ900439 |
| *Lcor21* | MitoCatch | 0.1 | YES | PZ900440 |
| *Lcor22* | MitoCatch | 0.1 | YES | PZ900441 |
| *Lcor23* | MitoCatch | 0.1 | YES | PZ900442 |
| *Lcor24* | MitoCatch | 0.1 | YES | PZ900443 |

(continue…)

| ***Sample*** | ***Mitogenome Assembly tool*** | ***Subset coverage*** | ***Complete Mitochondria Assembled*** | ***NCBI Acession Number*** |
|---|---|---|---|---|
| *Lcor25* | NOVOPlasty (Dierckxsens et al. 2016) | - | NO | - |
| *Ltim1* | MitoCatch | 0.1 | YES | PZ900467 |
| *Ltim2* | MitoCatch | 0.1 | YES | PZ900468 |
| *Ltim3* | MitoCatch | 0.1 | YES | PZ900469 |
| *Ltim4* | MitoCatch | 0.3 | YES | PZ900470 |
| *Ltim5* | MitoCatch | 0.1 | YES | PZ900471 |
| *Ltim6* | MitoCatch | 0.3 | YES | PZ900472 |
| *Lgra1* | MitoCatch | 0.1 | YES | PZ900458 |
| *Lgra2* | MitoCatch | 0.1 | YES | PZ900459 |
| *Lgra3* | MitoCatch | 0.1 | YES | PZ900460 |
| *Lgra4* | MitoCatch | 0.1 | YES | PZ900461 |
| *Lgra5* | MitoCatch | 0.1 | YES | PZ900462 |
| *Lgra6* | MitoCatch | 0.1 | YES | PZ900463 |
| *Lgra7* | MitoCatch | 0.3 | YES | PZ900464 |
| *Lgra8* | MitoCatch | 0.1 | YES | PZ900465 |
| *Lgra9* | MitoCatch | 0.3 | YES | PZ900466 |
| *Lgra10* | MitoCatch | 0.1 | YES | PZ900457 |
| *Leur1* | NOVOPlasty (Dierckxsens et al. 2016) | - | NO | - |
| *Leur2* | MitoCatch | 0.3 | YES | PZ920792 |
| *Leur3* | MitoCatch | 0.5 | YES | PZ900451 |
| *Leur4* | MitoCatch | 0.3 | YES | PZ900452 |
| *Leur5* | MitoCatch | 0.3 | YES | PZ900453 |
| *Leur6* | NOVOPlasty (Dierckxsens et al. 2016) | - | YES | PZ900454 |
| *Leur7* | NOVOPlasty (Dierckxsens et al. 2016) | - | YES | PZ900455 |
| *Leur8* | MitoCatch | 0.5 | YES | PZ900456 |
| *Leur9* | NOVOPlasty (Dierckxsens et al. 2016) | - | NO | - |
| *Leur10* | NOVOPlasty (Dierckxsens et al. 2016) | - | NO | - |
| *Lame1* | MitoCatch | 0.1 | YES | PZ900405 |
| *Lame2* | MitoCatch | 0.1 | YES | Waiting accession |
| *Lame3* | MitoCatch | 0.1 | YES | Waiting accession |
| *Lame4* | MitoCatch | 0.1 | YES | Waiting accession |
| *Lame5* | MitoCatch | 0.1 | YES | Waiting accession |
| *Lame6* | MitoCatch | 0.1 | YES | Waiting accession |
| *Lame7* | MitoCatch | 0.1 | YES | Waiting accession |
| *Lame8* | MitoCatch | 0.3 | YES | Waiting accession |
| *Lcor Iron 125* | EAGER (Peltzer et al. 2016) | - | NO | PZ900473 |
| *Lcor Iron 127* | EAGER (Peltzer et al. 2016) | - | NO | PZ900474 |
| *Lcor Iron 129* | EAGER (Peltzer et al. 2016) | - | NO | PZ900475 |

*Table S3 – AIC for each BPP Model*

| ***Model*** | ***LogLik*** | ***K*** | ***AIC*** | ***DeltaAIC*** |
|---|---|---|---|---|
| *MSC-I proto cascor more* | -1594395.269 | 22 | 3188834.538 | 0 |
| *MSC-I proto cascor* | -1594533.7665 | 22 | 3189111.533 | 276.9 |
| *MSC-I proto cas* | -1594621.7095 | 17 | 3189277.419 | 442.8 |
| *MSC-M proto cas* | -1594637.1875 | 11 | 3189296.375 | 461.8 |
| *MSC-M proto cascor* | -1594668.7065 | 13 | 3189363.413 | 528.8 |
| *MSC-M* | -1594701.1635 | 9 | 3189420.327 | 585.7 |
| *MSC-I only cascor* | -1594716.2365 | 17 | 3189466.473 | 631.9 |
| *MSC-I proto cor* | -1594754.294 | 17 | 3189542.588 | 708.0 |
| *MSC-I only cas* | -1594831.321 | 12 | 3189686.642 | 852.1 |
| *MSC-I only cor* | -1594867.8305 | 12 | 3189759.661 | 925.1 |
| *MSC-I proto* | -1595069.441 | 12 | 3190162.882 | 1328.3 |
| *MSC* | -1595106.06 | 7 | 3190226.12 | 1391.5 |

*Table S4 – Best Model Parameters*

| ***Parameters*** | ***Mean*** | ***Median*** | ***HPD 2.5%*** | ***HPD 97.5%*** |
|---|---|---|---|---|
| *Ne Ltim* | 125.7k | 123.9k | 60.3k | 193.6k |
| *Ne Lcas* | 79.5k | 76k | 50.2k | 115.6k |
| *Ne Lcor* | 97.2k | 92.1k | 43.9k | 161k |
| *Ne Anc* | 415.8k | 415.5k | 380.6k | 450.8k |
| *Ne proto* | 91.8k | 91.8k | 86.1k | 97.7k |
| *Div Ltim proto* | 2.51 mya | 2.51 mya | 2.36 mya | 2.66 mya |
| *Intr Ltim proto* | 919 kya | 917 kya | 834 kya | 1.01 mya |
| *intr Ltim Lcas* | 19 kya | 19 kya | 11 kya | 28 kya |
| *Intr Ltim Lcor* | 16 kya | 16 kya | 6 kya | 25 kya |
| *div Lcas Lcor* | 64 kya | 64 kya | 57 kya | 72 kya |
| *Intr precent Ltim proto* | 24.28% | 24.17% | 19.19% | 29.56% |
| *Intr precent Ltim Lcas* | 1.08% | 1.07% | 0.74% | 1.43% |
| *Intr precent Ltim Lcor* | 0.26% | 0.26% | 0.12% | 0.42% |

*Table S5 – All model parameters*

| ***Parameters*** | ***I mean*** | ***I median*** | ***I 2.5*** | ***I 97.5*** | ***II mean*** | ***II median*** | ***II 2.5*** | ***II 97.5*** | ***III mean*** | ***III median*** | ***III 2.5*** | ***III 97.5*** | ***IV mean*** | ***IV median*** | ***IV 2.5*** | ***IV 97.5*** |
|---|---|---|---|---|---|---|---|---|---|---|---|---|---|---|---|---|
| *Ne Ltim* | 528k | 527.9k | 510.7k | 545.5k | 596.9k | 596.7k | 572.1k | 621.7k | 245.2k | 228.5k | 45k | 469.6k | 294.3k | 293.7k | 180.2k | 407.6k |
| *Ne Lcas* | 32.8k | 32.8k | 29.6k | 36.2k | 35.4k | 35.4k | 32.1k | 38.9k | 32.8k | 32.8k | 29.5k | 36.2k | 78.9k | 74.5k | 50.7k | 117.5k |
| *Ne Lcor* | 89.1k | 89k | 79.8k | 98.4k | 96.8k | 96.6k | 87k | 106.9k | 112.9k | 103.9k | 28.1k | 209.9k | 91.9k | 91.8k | 80.8k | 102.8k |
| *Ne Anc* | 474.8k | 474.6k | 446.3k | 503.9k | 404.1k | 403.8k | 369k | 439.7k | 478.7k | 478.5k | 449.8k | 507.5k | 473.9k | 473.8k | 444.6k | 502.7k |
| *Ne Proto* | 118.8k | 118.8k | 112.7k | 124.7k | 102.2k | 102.2k | 96.2k | 108.3k | 117k | 117k | 110.9k | 122.9k | 108.1k | 108.1k | 102.3k | 114.2k |
| *Div Anc* | 1.68 mya | 1.68 mya | 1.61 mya | 1.75 mya | 2.61 mya | 2.61 mya | 2.46 mya | 2.77 mya | 1.67 mya | 1.67 mya | 1.6 mya | 1.74 mya | 1.7 mya | 1.7 mya | 1.63 mya | 1.78 mya |
| *Div Proto* | 71 kya | 71 kya | 63 kya | 77 kya | 75 kya | 75 kya | 69 kya | 82 kya | 69 kya | 69 kya | 61 kya | 76 kya | 61 kya | 61 kya | 54 kya | 71 kya |
| *Intr Ltim Lcas* | NA | NA | NA | NA | NA | NA | NA | NA | NA | NA | NA | NA | 24 kya | 24 kya | 10 kya | 36 kya |
| *Intr % Ltim Lcas* | NA | NA | NA | NA | NA | NA | NA | NA | NA | NA | NA | NA | 0.96% | 0.95% | 0.59% | 1.36% |
| *Intr Ltim Lcor* | NA | NA | NA | NA | NA | NA | NA | NA | 16 kya | 10 kya | 0 kya | 54 kya | NA | NA | NA | NA |
| *Intr Ltim Lcor* | NA | NA | NA | NA | NA | NA | NA | NA | 16 kya | 10 kya | 0 kya | 54 kya | NA | NA | NA | NA |
| *Intr % Ltim Lcor* | NA | NA | NA | NA | NA | NA | NA | NA | 0.16% | 0.15% | 0.05% | 0.30% | NA | NA | NA | NA |
| *Intr Ltim Proto* | NA | NA | NA | NA | 940 kya | 939 kya | 858 kya | 1.02 mya | NA | NA | NA | NA | NA | NA | NA | NA |
| *Intr % Ltim Proto* | NA | NA | NA | NA | 27.66% | 27.60% | 22.53% | 32.96% | NA | NA | NA | NA | NA | NA | NA | NA |
| *Ne Ltim* | NA | NA | NA | NA | NA | NA | NA | NA | NA | NA | NA | NA | NA | NA | NA | NA |
| *Ne Lcas* | NA | NA | NA | NA | NA | NA | NA | NA | NA | NA | NA | NA | NA | NA | NA | NA |
| *Ne Lcor* | NA | NA | NA | NA | NA | NA | NA | NA | NA | NA | NA | NA | NA | NA | NA | NA |
| *Ne Anc* | NA | NA | NA | NA | NA | NA | NA | NA | NA | NA | NA | NA | NA | NA | NA | NA |
| *Ne Proto* | NA | NA | NA | NA | NA | NA | NA | NA | NA | NA | NA | NA | NA | NA | NA | NA |
| *Div proto* | NA | NA | NA | NA | NA | NA | NA | NA | NA | NA | NA | NA | NA | NA | NA | NA |
| *Intr % Ltim proto* | NA | NA | NA | NA | NA | NA | NA | NA | NA | NA | NA | NA | NA | NA | NA | NA |
| *theta:5* | NA | NA | NA | NA | NA | NA | NA | NA | NA | NA | NA | NA | NA | NA | NA | NA |
| *theta:6* | NA | NA | NA | NA | NA | NA | NA | NA | NA | NA | NA | NA | NA | NA | NA | NA |
| *theta:7* | NA | NA | NA | NA | NA | NA | NA | NA | NA | NA | NA | NA | NA | NA | NA | NA |
| *theta:9* | NA | NA | NA | NA | NA | NA | NA | NA | NA | NA | NA | NA | NA | NA | NA | NA |
| *tau:7* | NA | NA | NA | NA | NA | NA | NA | NA | NA | NA | NA | NA | NA | NA | NA | NA |
| *tau:9* | NA | NA | NA | NA | NA | NA | NA | NA | NA | NA | NA | NA | NA | NA | NA | NA |
| *Mig Ltim Proto* | NA | NA | NA | NA | NA | NA | NA | NA | NA | NA | NA | NA | NA | NA | NA | NA |
| *Mig Ltim Lcas* | NA | NA | NA | NA | NA | NA | NA | NA | NA | NA | NA | NA | NA | NA | NA | NA |
| *Mig Ltim Lcor* | NA | NA | NA | NA | NA | NA | NA | NA | NA | NA | NA | NA | NA | NA | NA | NA |

(continue…)

| ***Parameters*** | ***V mean*** | ***V median*** | ***V 2.5*** | ***V 97.5*** | ***VI mean*** | ***VI median*** | ***VI 2.5*** | ***VI 97.5*** | ***VII mean*** | ***VII median*** | ***VII 2.5*** | ***VII 97.5*** | ***VIII mean*** | ***VIII median*** | ***VIII 2.5*** | ***VIII 97.5*** |
|---|---|---|---|---|---|---|---|---|---|---|---|---|---|---|---|---|
| *Ne Ltim* | 106.5k | 100.3k | 15.8k | 208.3k | 289.1k | 291k | 174.9k | 393.7k | 530.6k | 530.5k | 513.9k | 547.8k | 536k | 535.9k | 518.9k | 553.5k |
| *Ne Lcas* | 36.1k | 36.1k | 32.6k | 39.6k | 75.4k | 73.5k | 53.8k | 100.4k | 35.3k | 35.3k | 32.1k | 38.8k | 34.2k | 34.1k | 30.9k | 37.3k |
| *Ne Lcor* | 114.6k | 104.5k | 39.9k | 212k | 97k | 95.2k | 67.4k | 128.5k | 95.5k | 95.4k | 86.2k | 106k | 98.7k | 98.5k | 88.4k | 108.8k |
| *Ne Anc* | 416.1k | 415.8k | 381.3k | 451.3k | 459.2k | 459k | 431.3k | 488.1k | 420.9k | 420.7k | 390.6k | 452.5k | 427.1k | 427k | 395.8k | 457.8k |
| *Ne Proto* | 98.2k | 98.2k | 92.3k | 104.4k | 101.6k | 101.5k | 95.4k | 107.6k | 98.4k | 98.4k | 92.5k | 104.4k | 96k | 95.9k | 89.9k | 102.1k |
| *Div Anc* | 2.52 mya | 2.51 mya | 2.37 mya | 2.65 mya | 1.76 mya | 1.76 mya | 1.69 mya | 1.84 mya | 2.13 mya | 2.13 mya | 2.01 mya | 2.25 mya | 2.06 mya | 2.06 mya | 1.94 mya | 2.17 mya |
| *Div Proto* | NA | NA | NA | NA | 59 kya | 59 kya | 51 kya | 67 kya | 74 kya | 74 kya | 67 kya | 81 kya | 76 kya | 76 kya | 70 kya | 84 kya |
| *Intr Ltim Lcas* | NA | NA | NA | NA | NA | NA | NA | NA | NA | NA | NA | NA | NA | NA | NA | NA |
| *Intr % Ltim Lcas* | NA | NA | NA | NA | 1.20% | 1.20% | 0.83% | 1.58% | NA | NA | NA | NA | NA | NA | NA | NA |
| *Intr Ltim Lcor* | 13 kya | 11 kya | 1 kya | 28 kya | 28 kya | 29 kya | 16 kya | 39 kya | NA | NA | NA | NA | NA | NA | NA | NA |
| *Intr Ltim Lcor* | 13 kya | 11 kya | 1 kya | 28 kya | 37 kya | 34 kya | 20 kya | 60 kya | NA | NA | NA | NA | NA | NA | NA | NA |
| *Intr % Ltim Lcor* | 0.14% | 0.13% | 0.04% | 0.24% | 0.38% | 0.37% | 0.19% | 0.58% | NA | NA | NA | NA | NA | NA | NA | NA |
| *Intr Ltim Proto* | 869 kya | 868 kya | 792 kya | 945 kya | NA | NA | NA | NA | NA | NA | NA | NA | NA | NA | NA | NA |
| *Intr % Ltim Proto* | 24.40% | 24.34% | 19.72% | 29.11% | NA | NA | NA | NA | NA | NA | NA | NA | NA | NA | NA | NA |
| *Ne Ltim* | NA | NA | NA | NA | NA | NA | NA | NA | NA | NA | NA | NA | NA | NA | NA | NA |
| *Ne Lcas* | NA | NA | NA | NA | NA | NA | NA | NA | NA | NA | NA | NA | NA | NA | NA | NA |
| *Ne Lcor* | NA | NA | NA | NA | NA | NA | NA | NA | NA | NA | NA | NA | NA | NA | NA | NA |
| *Ne Anc* | NA | NA | NA | NA | NA | NA | NA | NA | NA | NA | NA | NA | NA | NA | NA | NA |
| *Ne Proto* | NA | NA | NA | NA | NA | NA | NA | NA | NA | NA | NA | NA | NA | NA | NA | NA |
| *Div proto* | 75 kya | 75 kya | 68 kya | 82 kya | NA | NA | NA | NA | NA | NA | NA | NA | NA | NA | NA | NA |
| *Intr % Ltim proto* | NA | NA | NA | NA | NA | NA | NA | NA | NA | NA | NA | NA | NA | NA | NA | NA |
| *theta:5* | 342.1k | 342k | 317.2k | 367.3k | NA | NA | NA | NA | NA | NA | NA | NA | NA | NA | NA | NA |
| *theta:6* | 716.8k | 715.5k | 661.9k | 773.8k | NA | NA | NA | NA | NA | NA | NA | NA | NA | NA | NA | NA |
| *theta:7* | 250.4k | 246.9k | 179.7k | 326.4k | NA | NA | NA | NA | NA | NA | NA | NA | NA | NA | NA | NA |
| *theta:9* | 93.8k | 92.9k | 72.2k | 116.3k | NA | NA | NA | NA | NA | NA | NA | NA | NA | NA | NA | NA |
| *tau:7* | 869 kya | 868 kya | 792 kya | 945 kya | NA | NA | NA | NA | NA | NA | NA | NA | NA | NA | NA | NA |
| *tau:9* | 13 kya | 11 kya | 1 kya | 28 kya | NA | NA | NA | NA | NA | NA | NA | NA | NA | NA | NA | NA |
| *Mig Ltim Proto* | NA | NA | NA | NA | NA | NA | NA | NA | 0.013838 | 0.013813 | 0.0113 | 0.0164 | 0.010189 | 0.010153 | 0.00770 | 0.01266 |
| *Mig Ltim Lcas* | NA | NA | NA | NA | NA | NA | NA | NA | NA | NA | NA | NA | 0.003526 | 0.003502 | 0.00235 | 0.00476 |
| *Mig Ltim Lcor* | NA | NA | NA | NA | NA | NA | NA | NA | NA | NA | NA | NA | 0.004129 | 0.004061 | 0.00246 | 0.00592 |

(continue…)

| Parameters | IX mean | IX median | IX 2.5 | IX 97.5 | X mean | X median | X 2.5 | X 97.5 | XI mean | XI median | XI 2.5 | XI 97.5 | XII mean | XII median | XII 2.5 | XII 97.5 |
|---|---|---|---|---|---|---|---|---|---|---|---|---|---|---|---|---|
| *Ne Ltim* | 537.1k | 537.1k | 520.1k | 554.6k | 113.4k | 112.2k | 60.5k | 166.5k | NA | NA | NA | NA | NA | NA | NA | NA |
| *Ne Lcas* | 34k | 34k | 30.7k | 37.1k | 108.5k | 100.2k | 56.6k | 181.3k | NA | NA | NA | NA | NA | NA | NA | NA |
| *Ne Lcor* | 100.6k | 100.4k | 90.2k | 111.2k | 99.9k | 99.7k | 88.6k | 111.7k | NA | NA | NA | NA | NA | NA | NA | NA |
| *Ne Anc* | 423.3k | 423k | 392.2k | 453.3k | 423.4k | 423k | 387.6k | 458.9k | NA | NA | NA | NA | NA | NA | NA | NA |
| *Ne Proto* | 94.8k | 94.8k | 89.2k | 101.2k | 94.5k | 94.4k | 88.8k | 100.1k | NA | NA | NA | NA | NA | NA | NA | NA |
| *Div Anc* | 2.09 mya | 2.09 mya | 1.97 mya | 2.2 mya | 2.48 mya | 2.48 mya | 2.34 mya | 2.64 mya | 2.49 mya | 2.49 mya | 2.35 mya | 2.64 mya | 2.51 mya | 2.51 mya | 2.36 mya | 2.66 mya |
| *Div Proto* | 76 kya | 76 kya | 69 kya | 83 kya | 66 kya | 66 kya | 59 kya | 73 kya | NA | NA | NA | NA | NA | NA | NA | NA |
| *Intr Ltim Lcas* | NA | NA | NA | NA | 16 kya | 15 kya | 9 kya | 25 kya | 19 kya | 19 kya | 11 kya | 26 kya | 19 kya | 19 kya | 11 kya | 28 kya |
| *Intr % Ltim Lcas* | NA | NA | NA | NA | 0.86% | 0.85% | 0.60% | 1.15% | 1.06% | 1.06% | 0.75% | 1.39% | 1.08% | 1.07% | 0.74% | 1.43% |
| *Intr Ltim Lcor* | NA | NA | NA | NA | NA | NA | NA | NA | 21 kya | 21 kya | 12 kya | 33 kya | 16 kya | 16 kya | 6 kya | 25 kya |
| *Intr Ltim Lcor* | NA | NA | NA | NA | NA | NA | NA | NA | 21 kya | 21 kya | 12 kya | 33 kya | 16 kya | 16 kya | 6 kya | 25 kya |
| *Intr % Ltim Lcor* | NA | NA | NA | NA | NA | NA | NA | NA | 0.31% | 0.30% | 0.15% | 0.48% | 0.26% | 0.26% | 0.12% | 0.42% |
| *Intr Ltim Proto* | NA | NA | NA | NA | 904 kya | 904 kya | 819 kya | 983 kya | 917 kya | 917 kya | 831 kya | 1 mya | 919 kya | 917 kya | 834 kya | 1.01 mya |
| *Intr % Ltim Proto* | NA | NA | NA | NA | 24.60% | 24.52% | 19.44% | 29.85% | 23.72% | 23.66% | 18.68% | 28.87% | NA | NA | NA | NA |
| *Ne Ltim* | NA | NA | NA | NA | NA | NA | NA | NA | 127.4k | 126.9k | 78.2k | 178.2k | 125.7k | 123.9k | 60.3k | 193.6k |
| *Ne Lcas* | NA | NA | NA | NA | NA | NA | NA | NA | 78.2k | 75.4k | 50.4k | 112k | 79.5k | 76k | 50.2k | 115.6k |
| *Ne Lcor* | NA | NA | NA | NA | NA | NA | NA | NA | 102.9k | 98.9k | 57.9k | 154.5k | 97.2k | 92.1k | 43.9k | 161k |
| *Ne Anc* | NA | NA | NA | NA | NA | NA | NA | NA | 414.8k | 414.6k | 380.7k | 450.2k | 415.8k | 415.5k | 380.6k | 450.8k |
| *Ne Proto* | NA | NA | NA | NA | NA | NA | NA | NA | 90.8k | 90.7k | 85k | 96.7k | 91.8k | 91.8k | 86.1k | 97.7k |
| *Div proto* | NA | NA | NA | NA | NA | NA | NA | NA | NA | NA | NA | NA | 64 kya | 64 kya | 57 kya | 72 kya |
| *Intr % Ltim proto* | NA | NA | NA | NA | NA | NA | NA | NA | NA | NA | NA | NA | 24.28% | 24.17% | 19.19% | 29.56% |
| *theta:5* | NA | NA | NA | NA | NA | NA | NA | NA | NA | NA | NA | NA | NA | NA | NA | NA |
| *theta:6* | NA | NA | NA | NA | NA | NA | NA | NA | NA | NA | NA | NA | NA | NA | NA | NA |
| *theta:7* | NA | NA | NA | NA | NA | NA | NA | NA | NA | NA | NA | NA | NA | NA | NA | NA |
| *theta:9* | NA | NA | NA | NA | NA | NA | NA | NA | NA | NA | NA | NA | NA | NA | NA | NA |
| *tau:7* | NA | NA | NA | NA | NA | NA | NA | NA | NA | NA | NA | NA | NA | NA | NA | NA |
| *tau:9* | NA | NA | NA | NA | NA | NA | NA | NA | NA | NA | NA | NA | NA | NA | NA | NA |
| *Mig Ltim Proto* | 0.01073 | 0.010699 | 0.00841 | 0.01311 | NA | NA | NA | NA | NA | NA | NA | NA | NA | NA | NA | NA |
| *Mig Ltim Lcas* | 0.00357 | 0.003539 | 0.00239 | 0.00478 | NA | NA | NA | NA | NA | NA | NA | NA | NA | NA | NA | NA |
| *Mig Ltim Lcor* | NA | NA | NA | NA | NA | NA | NA | NA | NA | NA | NA | NA | NA | NA | NA | NA |

*Table S6 – D-Statistics*

| P1 | P2 | P3 | Dstatistic | Z-score | P-value | f4-ratio | clustering sensitive | clustering robust |
|---|---|---|---|---|---|---|---|---|
| *Lcor* | Lcas | Lgra | 0.159729 | 10.6 | 2.3e-16 | 0.009 | 2.3e-16 | 2.3e-16 |
| *Lcor* | Lcas | Lgra | 0.0409204 | 7.9 | 2.6e-15 | 0.0027 | 2.3e-16 | 5.1e-10 |
| *Lcor* | Lcas | Lgra | 0.0341961 | 10.7 | 2.3e-16 | 0.0025 | 2.3e-16 | 2.3E-016 |

*Table S7 – Cactus4 fragment statistics*

| chr | fragment count | mean length | median length | mean snp | median snp |
|---|---|---|---|---|---|
| *chr1* | 22364 | 1441,67018422464 | 832 | 64,3843677338579 | 38 |
| *chr10* | 16713 | 1250,3746185604 | 722 | 58,6356728295338 | 35 |
| *chr11* | 14229 | 1350,35912572914 | 768 | 61,3903998875535 | 36 |
| *chr12* | 13260 | 1371,59562594268 | 766 | 60,629713423831 | 35 |
| *chr13* | 13053 | 1302,64973569294 | 753 | 59,1886156439132 | 35 |
| *chr14* | 13428 | 1356,25134048257 | 748 | 60,0107983318439 | 35 |
| *chr15* | 10426 | 1425,57644350661 | 796 | 65,5377901400345 | 38 |
| *chr16* | 11101 | 1387,01477344383 | 792 | 64,3590667507431 | 37 |
| *chr17* | 9907 | 1293,31916826486 | 761 | 60,8361764409003 | 37 |
| *chr18* | 9427 | 1229,67370319295 | 727 | 56,7258937095576 | 34 |
| *chr19* | 8521 | 1184,34514728318 | 699 | 57,8947306654148 | 35 |
| *chr2* | 20916 | 1463,64467393383 | 814 | 66,8314687320711 | 39 |
| *chr20* | 7360 | 1301,93274456521 | 716 | 60,5714673913043 | 36 |
| *chr21* | 4848 | 1158,30899339934 | 638,5 | 57,5876650165016 | 35 |
| *chr22* | 5235 | 1285,96618911174 | 726 | 63,9497612225405 | 38 |
| *chr23* | 3898 | 1093,75423293996 | 640 | 56,9353514622883 | 35 |
| *chr3* | 22128 | 1403,99692697035 | 801 | 63,4471710050614 | 37 |
| *chr4* | 20922 | 1381,51290507599 | 782 | 62,7604435522416 | 37 |
| *chr5* | 21050 | 1346,92760095011 | 760,5 | 61,3174821852731 | 36 |
| *chr6* | 15001 | 1331,14592360509 | 752 | 62,8861409239384 | 37 |
| *chr7* | 16195 | 1382,90657610373 | 780 | 64,0674899660389 | 38 |
| *chr8* | 15034 | 1415,13243315152 | 797 | 66,6274444592257 | 39 |
| *chr9* | 13919 | 1345,10474890437 | 773 | 62,3684891155973 | 37 |
| *TOTAL* | 308935 | 1353,05682101412 | 767 | 62,2859565928107 | 37 |

### *Table S8 – Cactus12 fragments statistics*

| chr | fragment count | mean length | median length | mean snp | median snp |
|---|---|---|---|---|---|
| *chr1* | 4674 | 213,097133076593 | 129 | 14,2028241335044 | 9 |
| *chr2* | 4916 | 229,718877135882 | 142 | 15,2139951179821 | 10 |
| *chr3* | 4598 | 213,555458895171 | 129 | 14,0739451935624 | 9 |
| *chr4* | 4090 | 205,61173594132 | 123 | 13,7601466992665 | 9 |
| *chr5* | 3731 | 203,579201286518 | 113 | 13,2500670061645 | 8 |
| *chr6* | 3006 | 195,650698602794 | 119 | 13,7411842980705 | 9 |
| *chr7* | 3096 | 223,487403100775 | 122 | 14,3084625322997 | 9 |
| *chr8* | 3087 | 209,063816002591 | 126 | 14,6326530612244 | 10 |
| *chr9* | 2895 | 191,615889464594 | 112 | 13,2321243523316 | 9 |
| *chr10* | 3013 | 201,385330235645 | 117 | 13,2110852970461 | 8 |
| *chr11* | 2630 | 258,728517110266 | 132 | 14,4102661596958 | 9 |
| *chr12* | 2263 | 202,595227574016 | 118 | 13,0181175430844 | 8 |
| *chr13* | 2123 | 225,746113989637 | 112 | 12,8290155440414 | 8 |
| *chr14* | 2488 | 216,175241157556 | 118 | 13,3814308681672 | 9 |
| *chr15* | 2216 | 204,048285198555 | 126 | 14,1398916967509 | 9 |
| *chr16* | 2026 | 210,295162882527 | 131 | 14,0809476801579 | 9 |
| *chr17* | 2031 | 193,712456917774 | 109 | 12,9049729197439 | 8 |
| *chr18* | 1444 | 175,747229916897 | 95 | 11,7777008310249 | 8 |
| *chr19* | 1689 | 173,295441089402 | 86 | 11,2652457075192 | 7 |
| *chr20* | 1475 | 189,783050847457 | 109 | 12,8054237288135 | 8 |
| *chr21* | 914 | 147,741794310722 | 89 | 10,5284463894967 | 7 |
| *chr22* | 791 | 185,230088495575 | 97 | 13,0328697850821 | 8 |
| *chr23* | 574 | 144,175958188153 | 75,5 | 9,41811846689895 | 7 |
| *TOTAL* | 59770 | 207,680759578383 | 119 | 13,6307679437845 | 9 |

### *Table S9 – SNP Heatmap High Coverage*

| Sample | Species | Blue | Yellow | Red | Missing | Total |
|---|---|---|---|---|---|---|
| *Lcas1* | Lcas | 6839904 | 40423 | 69766 | 18526 | 6968619 |
| *Lcas2* | Lcas | 6842541 | 35024 | 53408 | 37646 | 6968619 |
| *Lcas3* | Lcas | 6785345 | 27512 | 45952 | 109810 | 6968619 |
| *Lcas4* | Lcas | 6773958 | 27655 | 45268 | 121738 | 6968619 |
| *Lcas5* | Lcas | 6808181 | 36816 | 55253 | 68369 | 6968619 |
| *Lcas6* | Lcas | 6833478 | 40733 | 67548 | 26860 | 6968619 |
| *Lcas7* | Lcas | 6843695 | 39303 | 58433 | 27188 | 6968619 |
| *Lcas8* | Lcas | 6837974 | 35134 | 52929 | 42582 | 6968619 |
| *Lcas9* | Lcas | 6843619 | 40238 | 54295 | 30467 | 6968619 |
| *Lcas10* | Lcas | 6844949 | 35279 | 49777 | 38614 | 6968619 |
| *Lcor1* | Lcor | 6760170 | 33948 | 153299 | 21202 | 6968619 |
| *Lcor2* | Lcor | 6730568 | 36259 | 151333 | 50459 | 6968619 |
| *Lcor3* | Lcor | 6721175 | 34058 | 154360 | 59026 | 6968619 |
| *Lcor4* | Lcor | 6727246 | 34535 | 159528 | 47310 | 6968619 |
| *Lcor5* | Lcor | 6709563 | 34957 | 174987 | 49112 | 6968619 |
| *Lcor6* | Lcor | 6739004 | 45247 | 158744 | 25624 | 6968619 |
| *Lcor7* | Lcor | 6680421 | 44125 | 156673 | 87400 | 6968619 |
| *Lcor8* | Lcor | 6742496 | 47007 | 158315 | 20801 | 6968619 |
| *Lcor9* | Lcor | 6726900 | 37767 | 153104 | 50848 | 6968619 |
| *Lcor10* | Lcor | 6728308 | 44413 | 157966 | 37932 | 6968619 |
| *total* | | 135519495 | 750433 | 2130938 | 971514 | 139372380 |
| *Fraction of active sites* | | 97,92 | 0,54 | 1,54 | | |

*Table S10 – Snp Heatmap All samples*

| Sample | Species | Blue | Yellow | Red | Missing | Total |
|---|---|---|---|---|---|---|
| *Lcas1* | Lcas | 6818464 | 43389 | 88240 | 18526 | 6968619 |
| *Lcas2* | Lcas | 6823179 | 37552 | 70242 | 37646 | 6968619 |
| *Lcas3* | Lcas | 6765279 | 32945 | 60585 | 109810 | 6968619 |
| *Lcas4* | Lcas | 6754380 | 32954 | 59547 | 121738 | 6968619 |
| *Lcas5* | Lcas | 6788925 | 41284 | 70041 | 68369 | 6968619 |
| *Lcas6* | Lcas | 6818754 | 40521 | 82484 | 26860 | 6968619 |
| *Lcas7* | Lcas | 6824344 | 38566 | 78521 | 27188 | 6968619 |
| *Lcas8* | Lcas | 6821578 | 32010 | 72449 | 42582 | 6968619 |
| *Lcas9* | Lcas | 6823500 | 40467 | 74185 | 30467 | 6968619 |
| *Lcas10* | Lcas | 6825599 | 38327 | 66079 | 38614 | 6968619 |
| *Lcor1* | Lcas | 6809636 | 20519 | 117262 | 21202 | 6968619 |
| *Lcor2* | Lcas | 6780038 | 23018 | 115104 | 50459 | 6968619 |
| *Lcor3* | Lcas | 6770472 | 20791 | 118330 | 59026 | 6968619 |
| *Lcor4* | Lcas | 6776196 | 21474 | 123639 | 47310 | 6968619 |
| *Lcor5* | Lcas | 6757925 | 22000 | 139582 | 49112 | 6968619 |
| *Lcor6* | Lcas | 6780214 | 34270 | 128511 | 25624 | 6968619 |
| *Lcor7* | Lcas | 6719436 | 34357 | 127426 | 87400 | 6968619 |
| *Lcor8* | Lcas | 6782024 | 36860 | 128934 | 20801 | 6968619 |
| *Lcor9* | Lcas | 6768356 | 25813 | 123602 | 50848 | 6968619 |
| *Lcor10* | Lcas | 6768886 | 33309 | 128492 | 37932 | 6968619 |
| *Lcas11* | Lcas | 5227669 | 26745 | 51572 | 1662633 | 6968619 |
| *Lcas12* | Lcas | 5632142 | 28416 | 62395 | 1245666 | 6968619 |
| *Lcas13* | Lcor | 4841006 | 27853 | 50496 | 2049264 | 6968619 |
| *Lcas14* | Lcor | 5209801 | 28771 | 56154 | 1673893 | 6968619 |
| *Lcas15* | Lcor | 4891534 | 29659 | 51222 | 1996204 | 6968619 |
| *Lcas16* | Lcor | 5170216 | 26179 | 45056 | 1727168 | 6968619 |
| *Lcas17* | Lcor | 5894733 | 29843 | 50797 | 993246 | 6968619 |
| *Lcas18* | Lcor | 6218238 | 40194 | 66685 | 643502 | 6968619 |
| *Lcas19* | Lcor | 6274542 | 33475 | 57823 | 602779 | 6968619 |
| *Lcas20* | Lcor | 6263754 | 30982 | 54226 | 619657 | 6968619 |
| *Lcas21* | Lcor | 5091264 | 26830 | 47249 | 1803276 | 6968619 |
| *Lcas22* | Lcor | 5478363 | 27652 | 48736 | 1413868 | 6968619 |
| *Lcor11* | Lcor | 4993447 | 25866 | 86803 | 1862503 | 6968619 |
| *Lcor12* | Lcor | 4839718 | 23507 | 79333 | 2026061 | 6968619 |
| *Lcor13* | Lcor | 4627253 | 24223 | 80330 | 2236813 | 6968619 |
| *Lcor14* | Lcor | 4239542 | 22136 | 71642 | 2635299 | 6968619 |
| *Lcor15* | Lcor | 5317856 | 26632 | 91204 | 1532927 | 6968619 |
| *Lcor16* | Lcor | 5044496 | 27314 | 92778 | 1804031 | 6968619 |
| *Lcor17* | Lcor | 4571422 | 23500 | 76969 | 2296728 | 6968619 |
| *Lcor18* | Lcor | 4140129 | 20733 | 67724 | 2740033 | 6968619 |
| *Lcor19* | Lcor | 4529768 | 24342 | 80970 | 2333539 | 6968619 |
| *Lcor22* | Lcor | 4139723 | 22240 | 70358 | 2736298 | 6968619 |
| *Lcor23* | Lcor | 5081163 | 26754 | 87155 | 1773547 | 6968619 |
| *Lcor24* | Lcor | 5026660 | 25525 | 86153 | 1830281 | 6968619 |
| *Lcor25* | Lcor | 4693816 | 24055 | 79130 | 2171618 | 6968619 |
| *Total* | | 272322604 | 1371886 | 3830859 | 49999744 | 327525093 |
| *Fraction of active sites* | | 98,13 | 0,49 | 1,38 | | |

*Table S11 – Description of genes in longest Cactus regions.*

| ***Gene*** | ***Function*** |
|---|---|
| *ACAD11* | Initiates long-chain fatty acid β-oxidation in mitochondria and peroxisomes. |
| *ACKR4* | Chemokine-scavenging receptor that regulates immune cell migration by internalizing and degrading chemokines. |
| *TDRD1* | piRNA pathway protein involved in transposon silencing and maintenance of germline genome integrity during spermatogenesis. |
| *TNIK* | Serine/threonine kinase that activates Wnt and Hippo signaling to regulate transcription, cell proliferation, and development. |
| *TTC21B* | Ciliary transport protein involved in retrograde intraflagellar transport and regulation of Wnt/Hedgehog signaling. |
| *ZNF654* | Predicted zinc finger transcription factor involved in DNA binding and regulation of gene expression. |
| *ZNF3* | KRAB zinc finger transcription factor regulating RNA polymerase II-mediated gene expression |
| *ZNF26* | KRAB zinc finger transcription factor regulating DNA-templated transcription |
| *RNU6-1* | U6 spliceosomal RNA required for pre-mRNA splicing |
| *OR5V1* | G protein-coupled olfactory receptor mediating odor detection |
| *OR2G3* | G protein-coupled olfactory receptor mediating odor detection |
| *FCRL1* | B-cell surface receptor involved in immune signaling |
| *PDGFD* | Secreted growth factor promoting cell proliferation and migration |
| *USH2A* | Extracellular matrix protein required for retinal and inner ear function |
| *LDAH* | Lipid droplet-associated enzyme regulating cholesterol and triglyceride homeostasis |
| *ZNF717* | KRAB zinc finger transcriptional repressor |
| *ZFP2* | Krüppel zinc finger transcription factor regulating gene expression |
| *SPATA31D4* | Membrane-associated protein involved in spermatogenesis |
| *SPATA31D1* | Membrane-associated protein involved in spermatogenesis |
| *OR7A17* | G protein-coupled olfactory receptor mediating odor detection |
| *PRMT1* | Protein arginine methyltransferase regulating transcription, chromatin, and signaling |
| *OR52M1* | G protein-coupled olfactory receptor involved in odor detection and olfactory signal transduction |

*Table S12 – Top 0.1% largest void regions.*

| Chr | StrPos | EndPos | Size |
|---|---|---|---|
| chr19 | 30052400 | 30499755 | 447355 |
| chr10 | 75217934 | 75573611 | 355677 |
| chr11 | 98551695 | 98892317 | 340622 |
| chr19 | 27186121 | 27522650 | 336529 |
| chr2 | 176180972 | 176462281 | 281309 |
| chr21 | 37784559 | 38061563 | 277004 |
| chr10 | 74453495 | 74724645 | 271150 |
| chr19 | 30981172 | 31242653 | 261481 |
| chr20 | 21488021 | 21743186 | 255165 |
| chr21 | 42865486 | 43112213 | 246727 |
| chr19 | 25617965 | 25851925 | 233960 |
| chr11 | 93032044 | 93265347 | 233303 |
| chr19 | 22662320 | 22880971 | 218651 |
| chr21 | 23454837 | 23669536 | 214699 |
| chr11 | 96230319 | 96444650 | 214331 |
| chr10 | 74999876 | 75214169 | 214293 |
| chr11 | 96792168 | 96998828 | 206660 |
| chr10 | 73967784 | 74170196 | 202412 |
| chr19 | 26071133 | 26269104 | 197971 |
| chr11 | 117302540 | 117480123 | 177583 |
| chr21 | 38826360 | 39003401 | 177041 |
| chr21 | 36262120 | 36432089 | 169969 |
| chr21 | 40074964 | 40242784 | 167820 |
| chr11 | 92617986 | 92782981 | 164995 |
| chr11 | 95575866 | 95740261 | 164395 |
| chr19 | 29271163 | 29434262 | 163099 |
| chr19 | 26501754 | 26663068 | 161314 |
| chr11 | 102087652 | 102247994 | 160342 |
| chr19 | 31779168 | 31935313 | 156145 |
| chr21 | 38672108 | 38824631 | 152523 |
| chr11 | 95807105 | 95948824 | 141719 |
| chr3 | 27672161 | 27812801 | 140640 |
| chr21 | 33444300 | 33583085 | 138785 |
| chr21 | 40779292 | 40917295 | 138003 |
| chr11 | 95066025 | 95201203 | 135178 |
| chr19 | 29120502 | 29252317 | 131815 |
| chr7 | 130764806 | 130894741 | 129935 |
| chr11 | 99631570 | 99760930 | 129360 |
| chr19 | 25443413 | 25572637 | 129224 |
| chr19 | 26374742 | 26501651 | 126909 |
| chr5 | 145053897 | 145180510 | 126613 |
| chr11 | 100597021 | 100722241 | 125220 |
| chr21 | 42631754 | 42756546 | 124792 |
| chr19 | 26888567 | 27013167 | 124600 |
| chr21 | 34853001 | 34975417 | 122416 |
| chr9 | 52677493 | 52797345 | 119852 |
| chr21 | 34362869 | 34479466 | 116597 |
| chr11 | 92912230 | 93026459 | 114229 |
| chr11 | 92493810 | 92607566 | 113756 |

(continue…)

| *Chr* | *StrPos* | *EndPos* | *Size* |
|---|---|---|---|
| *chr10* | 74724667 | 74837424 | 112757 |
| *chr19* | 27530639 | 27641975 | 111336 |
| *chr21* | 46176453 | 46287205 | 110752 |
| *chr19* | 27807975 | 27918299 | 110324 |
| *chr19* | 24031264 | 24137373 | 106109 |
| *chr19* | 22061146 | 22165344 | 104198 |
| *chr6* | 84137881 | 84241873 | 103992 |
| *chr19* | 24549762 | 24652911 | 103149 |
| *chr18* | 65519171 | 65620772 | 101601 |
| *chr7* | 132110152 | 132211395 | 101243 |
| *chr11* | 103324057 | 103425188 | 101131 |
| *chr6* | 66460542 | 66561309 | 100767 |
| *chr19* | 24675770 | 24775686 | 99916 |
| *chr19* | 27072622 | 27172119 | 99497 |
| *chr21* | 41857732 | 41956164 | 98432 |
| *chr19* | 29510448 | 29608428 | 97980 |
| *chr21* | 29182192 | 29280126 | 97934 |
| *chr11* | 102248053 | 102345956 | 97903 |
| *chr21* | 23919407 | 24017084 | 97677 |
| *chr21* | 32802086 | 32899720 | 97634 |
| *chr21* | 44297712 | 44395318 | 97606 |
| *chr21* | 45307566 | 45405128 | 97562 |
| *chr12* | 104921210 | 105017818 | 96608 |
| *chr19* | 28623218 | 28718866 | 95648 |
| *chr6* | 82915876 | 83011036 | 95160 |
| *chr19* | 30640221 | 30735275 | 95054 |
| *chr11* | 96118774 | 96210081 | 91307 |
| *chr11* | 97409660 | 97500759 | 91099 |
| *chr11* | 99761044 | 99851946 | 90902 |
| *chr9* | 51898164 | 51988990 | 90826 |
| *chr19* | 25103480 | 25192537 | 89057 |
| *chr21* | 35182445 | 35270567 | 88122 |
| *chr21* | 41385487 | 41473441 | 87954 |
| *chr19* | 21801413 | 21889357 | 87944 |
| *chr21* | 41981603 | 42069391 | 87788 |
| *chr20* | 21743190 | 21830926 | 87736 |
| *chr19* | 28043148 | 28130658 | 87510 |
| *chr11* | 96623288 | 96709637 | 86349 |
| *chr11* | 99478061 | 99564400 | 86339 |
| *chr21* | 21637356 | 21723393 | 86037 |
| *chr5* | 144430268 | 144515424 | 85156 |
| *chr19* | 24415250 | 24500139 | 84889 |
| *chr19* | 29720946 | 29805810 | 84864 |
| *chr21* | 37056429 | 37141058 | 84629 |
| *chr11* | 95224464 | 95308421 | 83957 |
| *chr8* | 109040616 | 109124381 | 83765 |
| *chr21* | 42779031 | 42862787 | 83756 |
| *chr22* | 43860747 | 43944026 | 83279 |
| *chr21* | 39400373 | 39483589 | 83216 |
| *chr21* | 41220088 | 41302733 | 82645 |
| *chr18* | 73758685 | 73839843 | 81158 |
| *chr11* | 94903178 | 94983963 | 80785 |

(continue…)

| *Chr* | *StrPos* | *EndPos* | *Size* |
|---|---|---|---|
| *chr12* | 90380082 | 90460691 | 80609 |
| *chr11* | 99048461 | 99128048 | 79587 |
| *chr21* | 41562031 | 41641305 | 79274 |
| *chr19* | 30554212 | 30633452 | 79240 |
| *chr12* | 101070162 | 101148855 | 78693 |
| *chr11* | 61910393 | 61988780 | 78387 |
| *chr5* | 159191770 | 159269725 | 77955 |
| *chr19* | 24788589 | 24866256 | 77667 |
| *chr11* | 94685853 | 94763279 | 77426 |
| *chr5* | 144736778 | 144813310 | 76532 |
| *chr18* | 76304583 | 76381089 | 76506 |
| *chr6* | 81707426 | 81783759 | 76333 |
| *chr14* | 80193126 | 80268691 | 75565 |
| *chr19* | 25970508 | 26045240 | 74732 |
| *chr8* | 107281725 | 107355940 | 74215 |
| *chr21* | 60484610 | 60558799 | 74189 |
| *chr17* | 80475750 | 80549604 | 73854 |
| *chr14* | 90453831 | 90527552 | 73721 |
| *chr21* | 39948897 | 40022447 | 73550 |
| *chr19* | 30810226 | 30883262 | 73036 |
| *chr2* | 48405463 | 48478314 | 72851 |
| *chr21* | 58245135 | 58317618 | 72483 |
| *chr11* | 94021137 | 94093247 | 72110 |
| *chr21* | 31868958 | 31940571 | 71613 |
| *chr18* | 71466482 | 71537891 | 71409 |
| *chr19* | 27968426 | 28039160 | 70734 |
| *chr21* | 37361139 | 37431789 | 70650 |
| *chr21* | 40477127 | 40547470 | 70343 |
| *chr2* | 48153095 | 48222867 | 69772 |
| *chr7* | 129325847 | 129395333 | 69486 |
| *chr21* | 35373398 | 35442865 | 69467 |
| *chr7* | 129923313 | 129992726 | 69413 |
| *chr8* | 10902832 | 10972235 | 69403 |
| *chr20* | 13496744 | 13566078 | 69334 |
| *chr12* | 100556895 | 100625853 | 68958 |
| *chr23* | 3169564 | 3237932 | 68368 |
| *chr6* | 82215740 | 82283725 | 67985 |
| *chr19* | 29615963 | 29683851 | 67888 |
| *chr21* | 44958362 | 45026207 | 67845 |
| *chr18* | 69570023 | 69637487 | 67464 |
| *chr18* | 64801093 | 64868108 | 67015 |
| *chr21* | 21733626 | 21800472 | 66846 |
| *chr17* | 83383580 | 83450300 | 66720 |
| *chr10* | 75763440 | 75829935 | 66495 |
| *chr20* | 34691109 | 34757590 | 66481 |
| *chr11* | 116992269 | 117058684 | 66415 |
| *chr19* | 23948784 | 24015082 | 66298 |
| *chr21* | 46438527 | 46504520 | 65993 |
| *chr4* | 101911675 | 101977662 | 65987 |
| *chr19* | 25037499 | 25103468 | 65969 |
| *chr21* | 41662387 | 41727921 | 65534 |
| *chr2* | 170938342 | 171003691 | 65349 |

(continue…)

| *Chr* | *StrPos* | *EndPos* | *Size* |
|---|---|---|---|
| *chr7* | 129173620 | 129238756 | 65136 |
| *chr19* | 31372668 | 31437449 | 64781 |
| *chr9* | 15597157 | 15661929 | 64772 |
| *chr19* | 31442480 | 31506737 | 64257 |
| *chr2* | 46644167 | 46708369 | 64202 |
| *chr21* | 29571234 | 29635243 | 64009 |
| *chr11* | 100062429 | 100126340 | 63911 |
| *chr12* | 100880415 | 100943965 | 63550 |
| *chr8* | 15061433 | 15124779 | 63346 |
| *chr11* | 116712172 | 116775351 | 63179 |
| *chr2* | 46498665 | 46561432 | 62767 |
| *chr21* | 41308021 | 41370734 | 62713 |
| *chr21* | 42562746 | 42625305 | 62559 |
| *chr18* | 68015426 | 68077877 | 62451 |
| *chr21* | 21801974 | 21864329 | 62355 |
| *chr21* | 29636272 | 29698492 | 62220 |
| *chr10* | 75849371 | 75911474 | 62103 |
| *chr2* | 47058102 | 47120039 | 61937 |
| *chr8* | 9347926 | 9409671 | 61745 |
| *chr11* | 94984001 | 95045718 | 61717 |
| *chr5* | 145585441 | 145646918 | 61477 |
| *chr20* | 8315394 | 8376591 | 61197 |
| *chr21* | 45107591 | 45168762 | 61171 |
| *chr4* | 165802597 | 165863373 | 60776 |
| *chr21* | 39263603 | 39324230 | 60627 |
| *chr19* | 25909926 | 25970507 | 60581 |
| *chr10* | 105776947 | 105837469 | 60522 |
| *chr2* | 2999882 | 3060219 | 60337 |
| *chr21* | 35280919 | 35341180 | 60261 |
| *chr21* | 34134195 | 34194422 | 60227 |
| *chr21* | 55663912 | 55723978 | 60066 |
| *chr5* | 139206010 | 139265493 | 59483 |
| *chr11* | 98211735 | 98271181 | 59446 |
| *chr5* | 144854046 | 144913485 | 59439 |
| *chr21* | 24122971 | 24182376 | 59405 |
| *chr21* | 36051957 | 36111249 | 59292 |
| *chr19* | 24173243 | 24232245 | 59002 |
| *chr20* | 44943101 | 45002103 | 59002 |
| *chr17* | 84238235 | 84297055 | 58820 |
| *chr10* | 75609336 | 75668097 | 58761 |
| *chr21* | 39700562 | 39759183 | 58621 |
| *chr5* | 151736797 | 151795022 | 58225 |
| *chr11* | 97877300 | 97935185 | 57885 |
| *chr11* | 98968657 | 99026502 | 57845 |
| *chr21* | 30693166 | 30750982 | 57816 |
| *chr8* | 475676 | 533320 | 57644 |
| *chr8* | 173748 | 231368 | 57620 |
| *chr11* | 93572145 | 93629729 | 57584 |
| *chr18* | 70909954 | 70967499 | 57545 |
| *chr21* | 37521059 | 37578341 | 57282 |
| *chr8* | 63021 | 119907 | 56886 |
| *chr23* | 25483092 | 25539958 | 56866 |

(continue…)

| ***Chr*** | ***StrPos*** | ***EndPos*** | ***Size*** |
|---|---|---|---|
| *chr19* | 23852794 | 23909603 | 56809 |
| *chr9* | 17214903 | 17271697 | 56794 |
| *chr8* | 10840375 | 10897125 | 56750 |
| *chr8* | 13688613 | 13745317 | 56704 |
| *chr19* | 24902732 | 24959266 | 56534 |
| *chr18* | 76213747 | 76270180 | 56433 |
| *chr13* | 96292029 | 96348259 | 56230 |
| *chr21* | 38449464 | 38505678 | 56214 |
| *chr11* | 92794519 | 92850462 | 55943 |
| *chr21* | 41503000 | 41558834 | 55834 |
| *chr3* | 32995321 | 33050646 | 55325 |
| *chr21* | 46073892 | 46129168 | 55276 |
| *chr6* | 81920336 | 81975558 | 55222 |
| *chr14* | 52869300 | 52924259 | 54959 |

***Supplementary Figures***

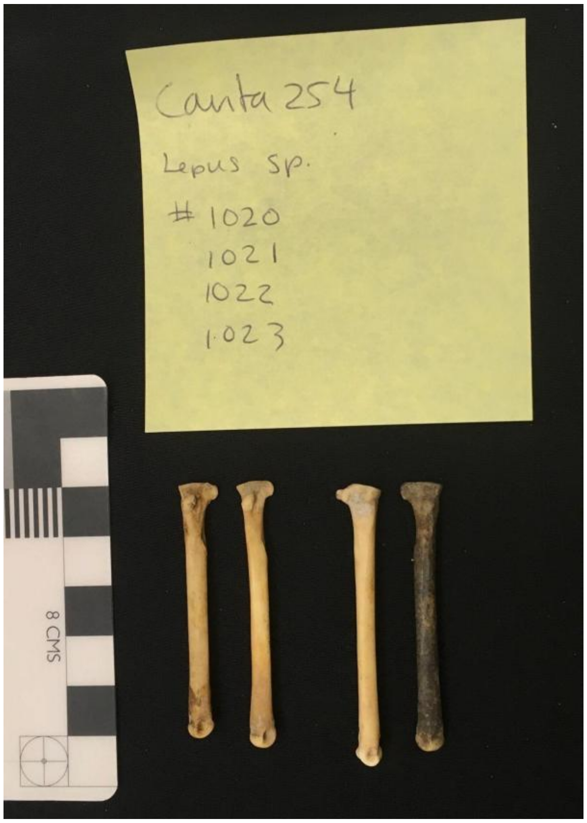


**Figure S1. Iron Age sampling.** Photographs of the *Lepus corsicanus* bone samples from Cavità 254 (Orvieto, Italy) used for ancient DNA analysis in this study.

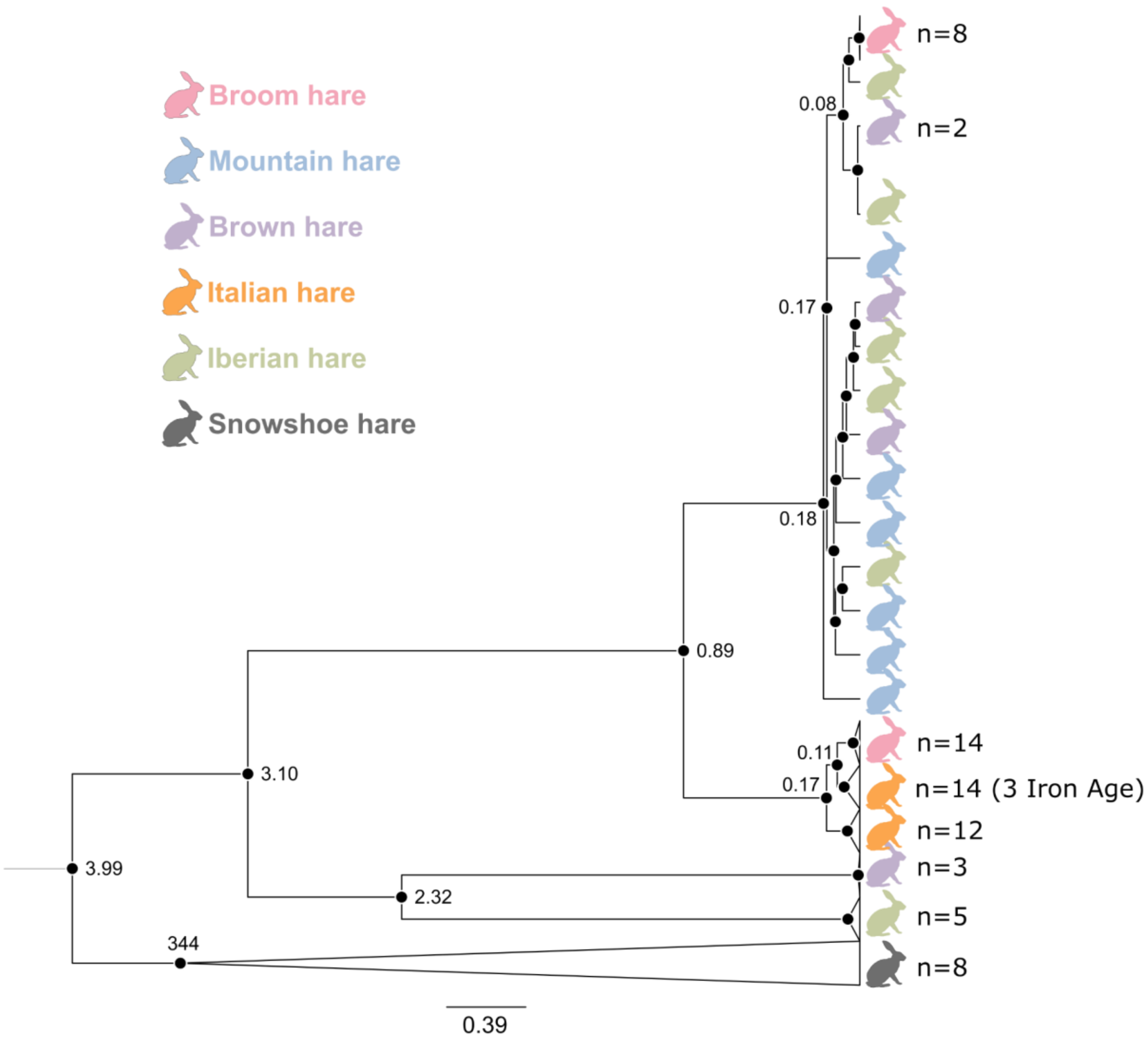


**Supplementary Figure 2. Mitogenome phylogeny.** Branch tips are color-coded by species according to the legend. Node labels indicate divergence times (Mya) followed by bootstrap support. Collapsed clades are shown as triangles, with sample sizes indicated.

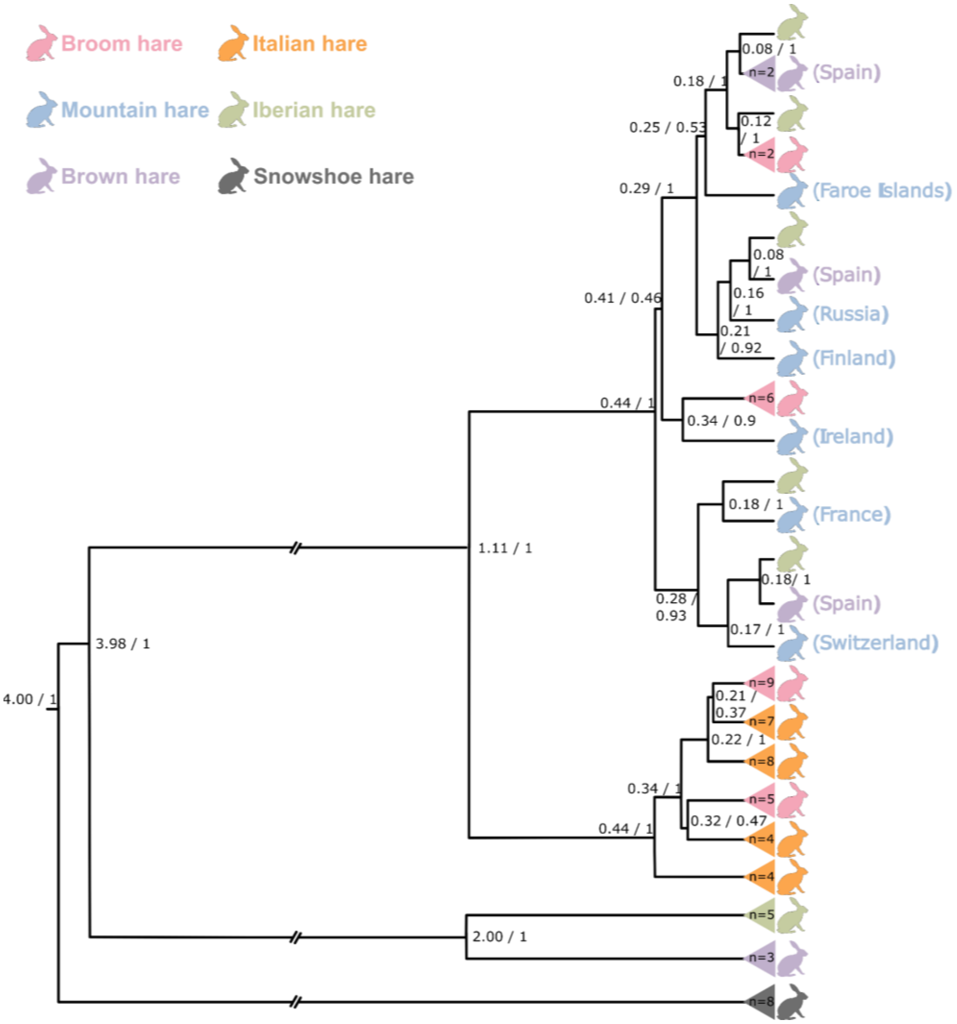


**Figure S3. Mitogenome phylogeny.** Branch tips are color-coded by species according to the legend. Node labels indicate divergence times (Mya) followed by bootstrap support. Collapsed clades are shown as colored triangles, with sample sizes indicated within each triangle. For species with broad geographic distributions (brown and mountain hare), sampling locations are given in parentheses next to sample names.

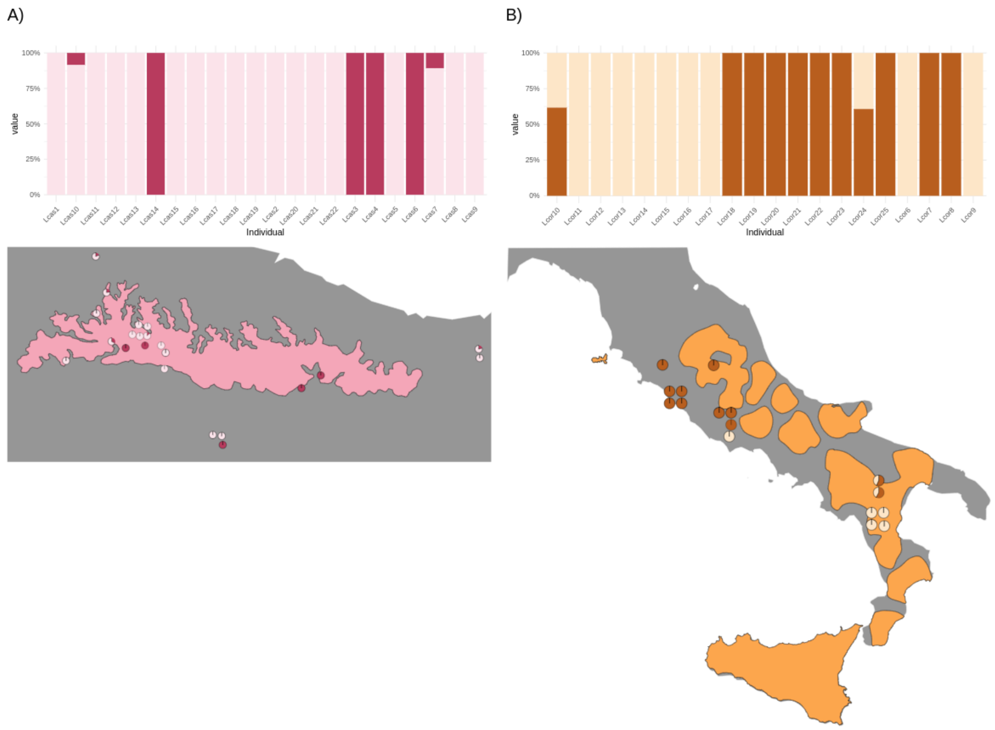


**Figure S4. Admixture plots for the best K (K=2) for the broom hare (top left) and the Italian hare (top right).** Map of the supposed distribution of the broom hare (bottom left) and Italian hare (bottom right) with pie charts representing the individual membership proportion obtained for each cluster.

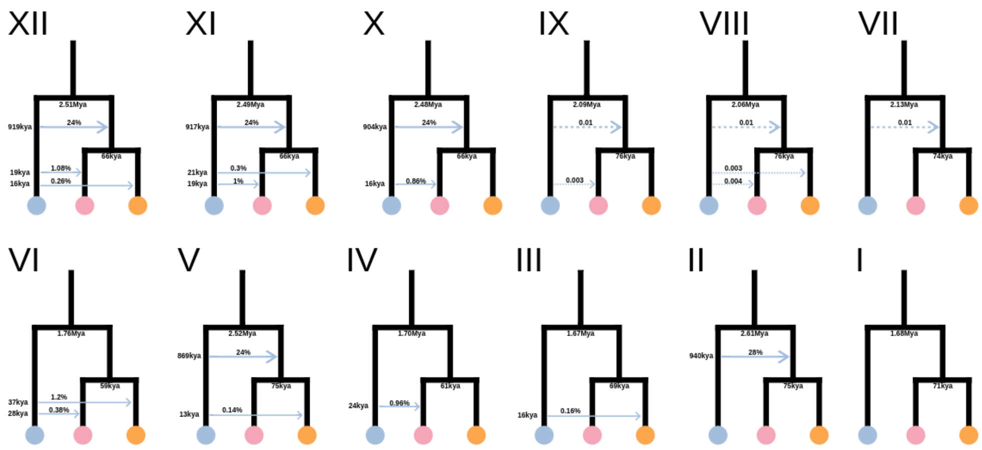


**Figure S5. Schematic representation of all demographic models tested with BPP.** Solid arrows indicate episodic introgression events, whereas dashed arrows indicate continuous migration. Divergence and introgression times are annotated in million years ago (Mya). Numbers above arrows represent the proportion of lineages affected in introgression models or the proportion of migrants per generation in migration models.

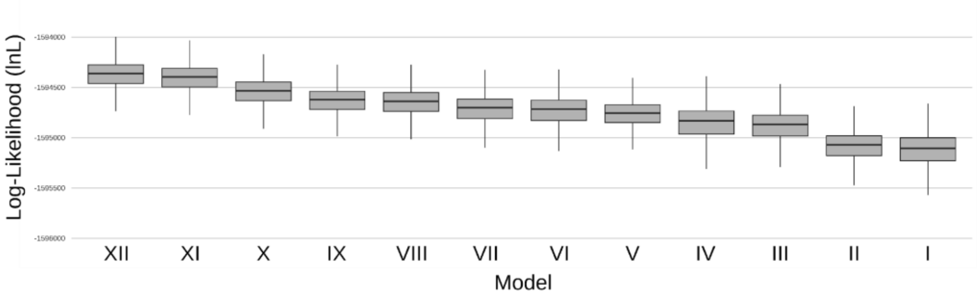


**Figure S6. Log-likelihood distributions for all tested BPP demographic models (described in Fig. S3).**

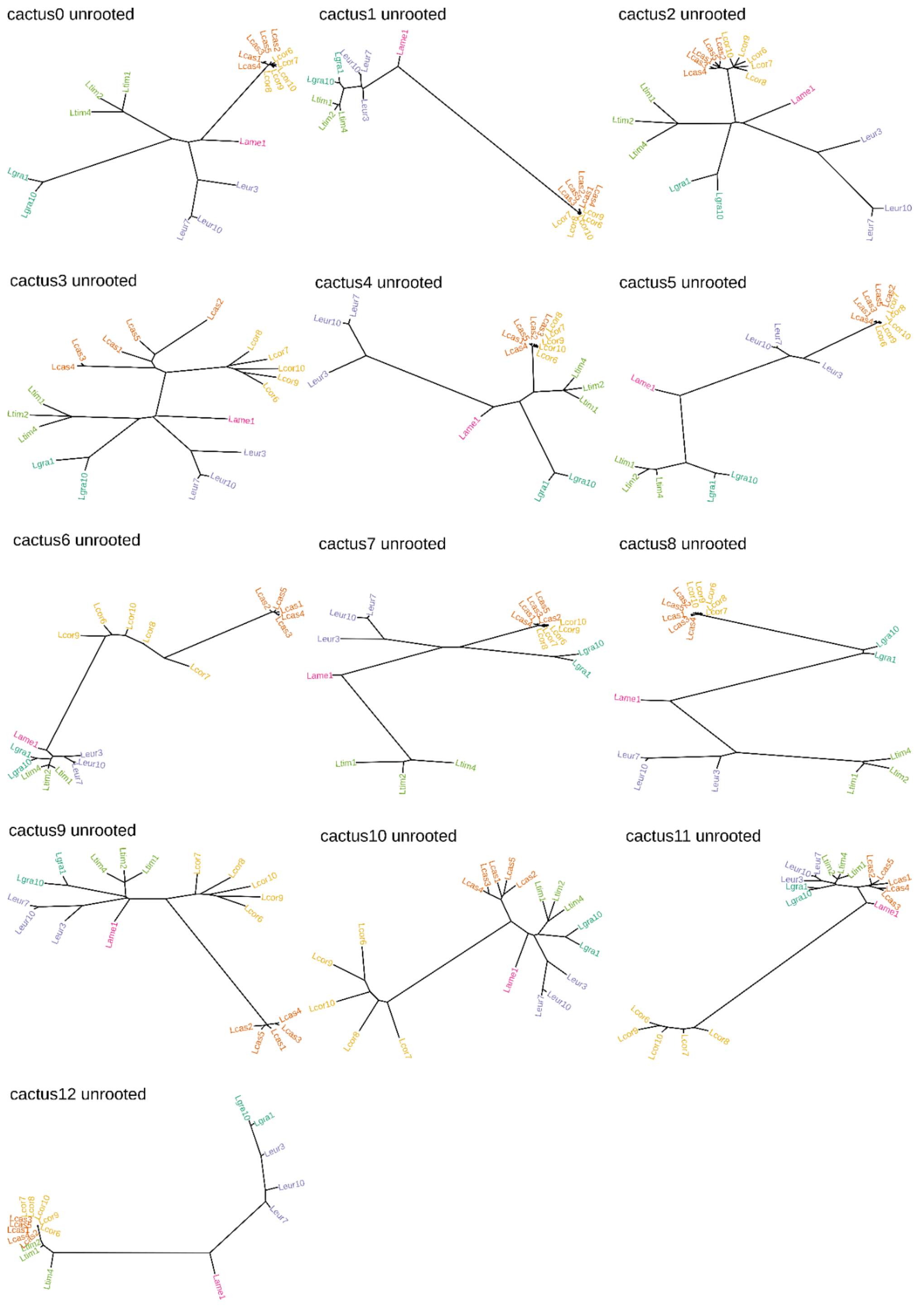


**Figure S7. Phylogenetic patterns ("cacti") identified by Saguaro across the genome.** All cacti inferred by Saguaro are shown as unrooted phylogenies representing distinct local genomic relationships among hare samples. Sample IDs at the tips correspond to those described in table S1 and are coloured according to species as in Fig. 1 and S1.

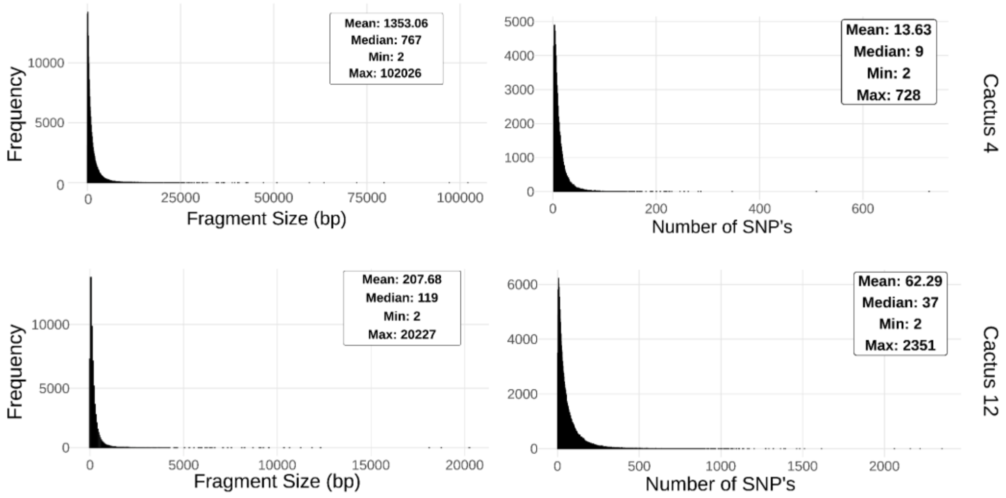


**Figure S8. Distribution of fragment size and number of SNPs for cactus 4 (top) and cactus 12 (bottom).** Left: fragment size distribution. Right: distribution of the number of SNPs per fragment.

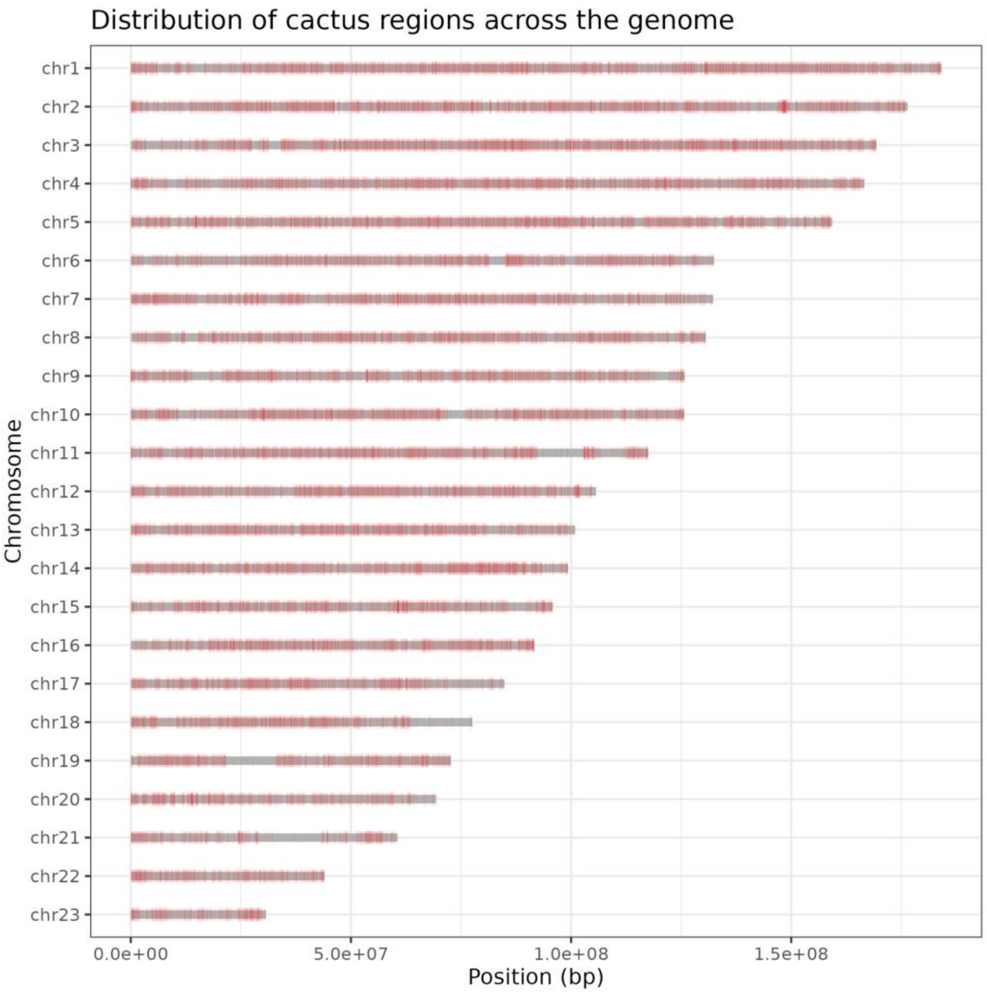


**Figure S9. Chromosomal distribution of cactus 4 and 12 segments (red bars).**

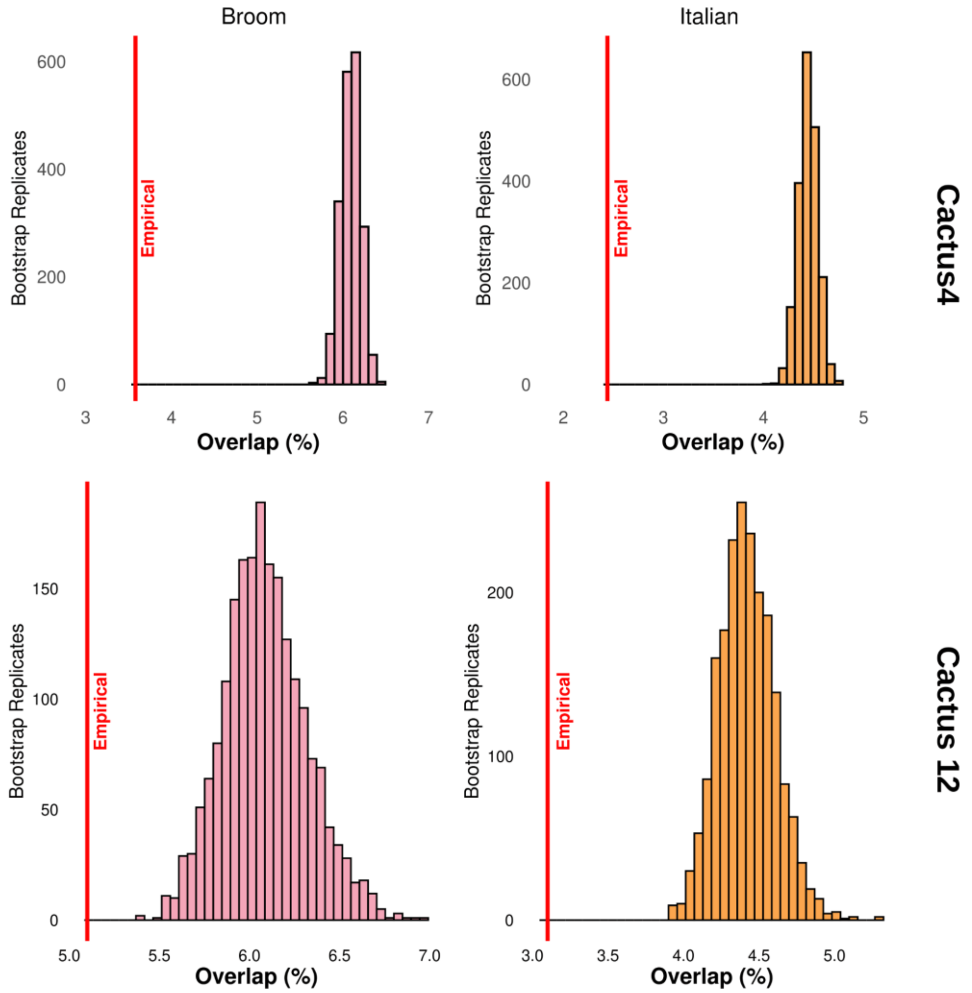


**Figure S10. Percentage overlap between low-mapping-quality regions identified by NGSparalog and circular bootstrap fragments for cactus 4 (top) and cactus 12 (bottom), for the broom hare (pink) and the Italian hare (orange).** Empirical values for each region set are indicated by a red line.

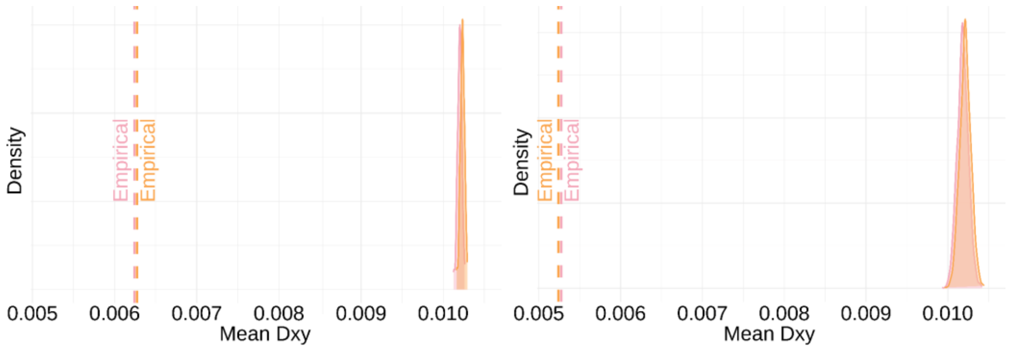


**Figure S11. Reduced sequence divergence between mountain and broom-Italian hares in cactus 4 and 12 segments.** Distribution of Dxy values obtained from circular bootstrap replicates for broom–mountain hare (pink) and Italian–mountain hare (orange) comparisons within cactus 4 (left) and cactus 12 (right). Dashed lines indicate the empirical Dxy values observed for each species pair within the corresponding cactus regions. Lower empirical Dxy values relative to the bootstrap distributions indicate reduced divergence consistent with introgression.

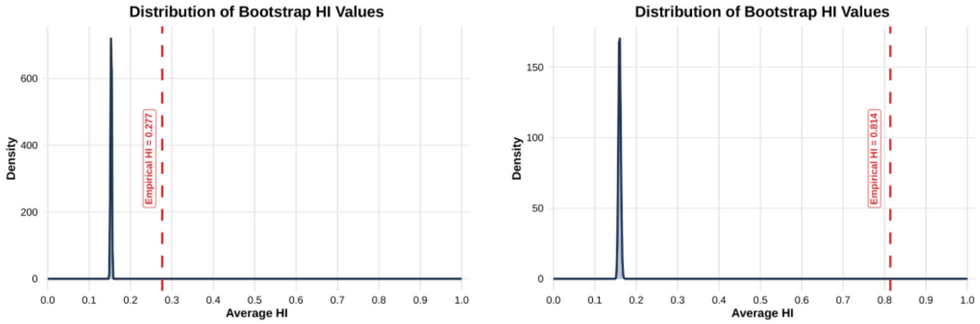


**Figure S12. Elevated mountain hare HI (Hybrid Index) in cactus 4 and 12 segments.** Distribution of mean mountain hare HI values obtained from circular bootstrap replicates for cactus 4 (left) and cactus 12 (right). The dashed red line indicates the empirical mean HI observed within each cactus region. Higher empirical HI values relative to the bootstrap distributions indicate enrichment for shared ancestry in the introgressed regions.

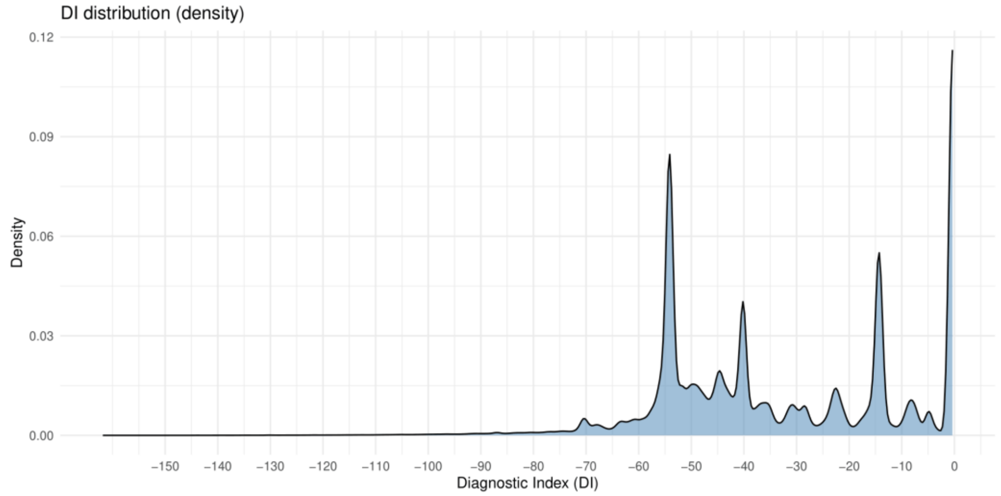


**Figure S13. Distribution of DI (diagnostic Index) values across sites.** Density plot showing the distribution of DI values. The y-axis represents density, and the x-axis shows DI values.

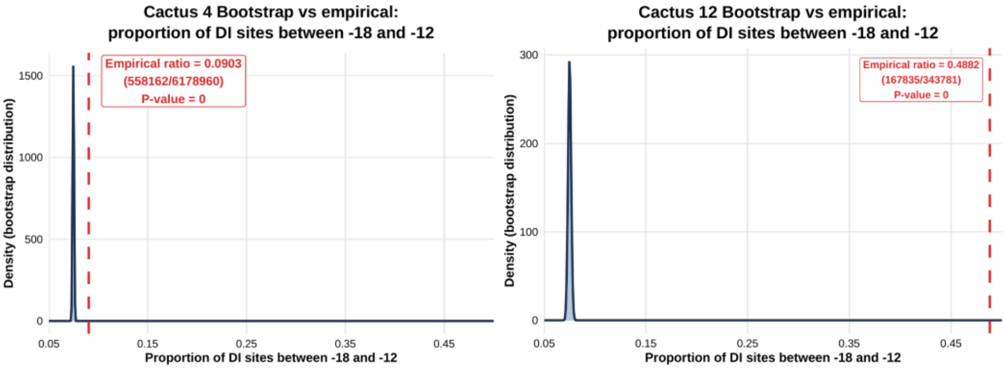


**Figure S14. Enrichment of elevated Diagnostic Index (DI) sites in cactus 4 and 12 segments.** Distribution of the proportion of elevated DI sites obtained from circular bootstrap replicates for cactus 4 (left) and cactus 12 (right). The dashed line indicates the empirical proportions.

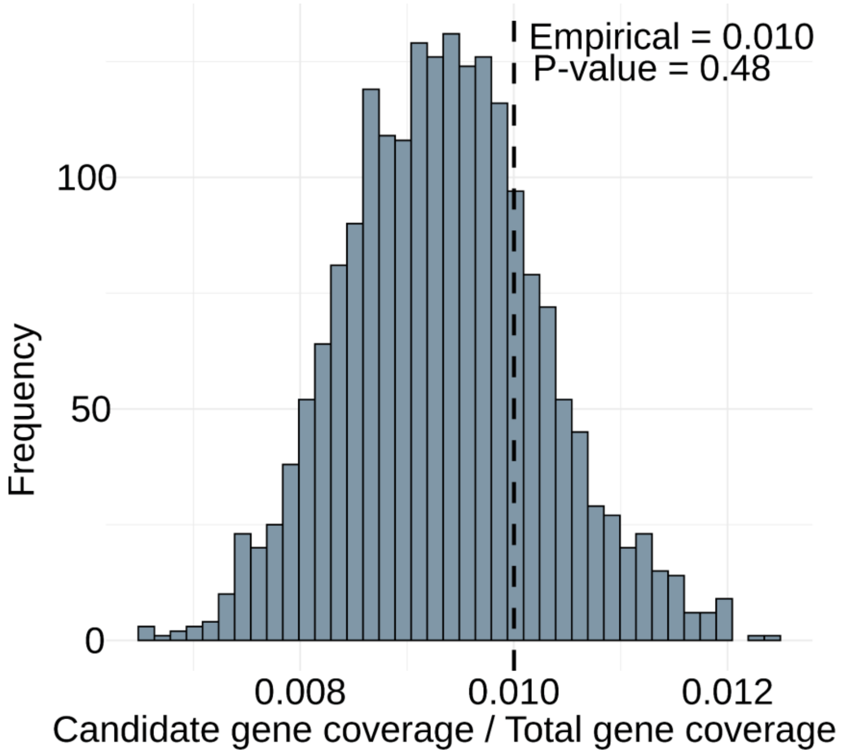


**Figure S15. Relative coverage of candidate genes within cactus 4 segments.** Histogram showing the distribution of the ratio between the number of base pairs overlapping candidate genes and the total number of base pairs overlapping annotated genes across 2,000 circular bootstrap replicates of cactus 4 segments. The vertical dashed line indicates the empirical value observed for cactus 4.

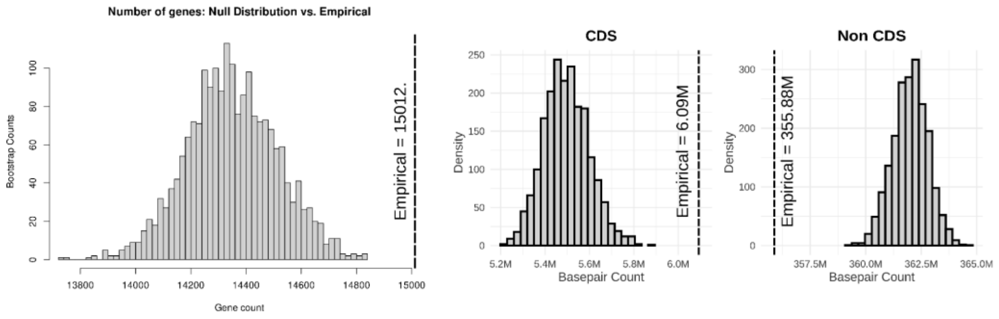


**Figure S16. Gene and coding sequence (CDS) content of cactus 4 compared with circular bootstrap replicates.** Left: distribution of the number of genes overlapping the 2,000 circular bootstrap replicates of cactus 4. The black dashed line indicates the empirical value. Right: distributions of the number of base pairs overlapping CDS and non-CDS regions across the 2,000 circular bootstrap replicates. Dashed lines indicate the corresponding empirical values for cactus 4.

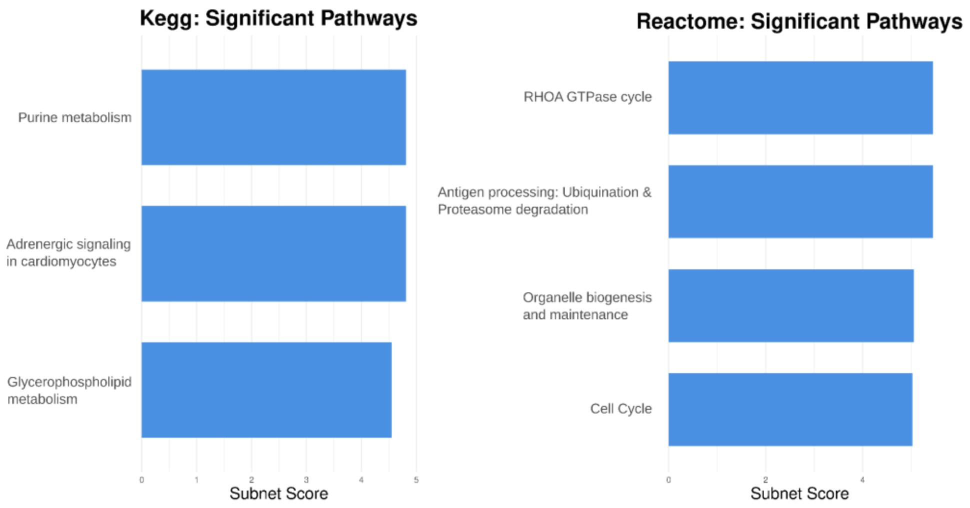


**Figure S17. SIGNET results for cactus 4 and 12 regions.** Significantly enriched pathways identified using the KEGG database (left) and the Reactome database (right).

### *References*


Best J, Doherty S, Armit I, Boev Z, Büster L, Cunliffe B, Foster A, Frimet B, Hamilton-Dyer S, Higham T, et al. 2022. Redefining the timing and circumstances of the chicken's introduction to Europe and north-west Africa. *Antiquity* 96:868–882.

Bouckaert R, Vaughan TG, Barido-Sottani J, Duchêne S, Fourment M, Gavryushkina A, Heled J, Jones G, Kühnert D, De Maio N, et al. 2019. BEAST 2.5: An advanced software platform for Bayesian evolutionary analysis.Pertea M, editor. *PLoS Comput Biol* 15:e1006650.

Carøe C, Gopalakrishnan S, Vinner L, Mak SST, Sinding MHS, Samaniego JA, Wales N, Sicheritz-Pontén T, Gilbert MTP. 2018. Single-tube library preparation for degraded DNA.Johnston S, editor. *Methods Ecol Evol* 9:410–419.

Dabney J, Knapp M, Glocke I, Gansauge M-T, Weihmann A, Nickel B, Valdiosera C, García N, Pääbo S, Arsuaga J-L, et al. 2013. Complete mitochondrial genome sequence of a Middle Pleistocene cave bear reconstructed from ultrashort DNA fragments. Proc. Natl. Acad. Sci. U.S.A. 110:15758–15763.

Damgaard PB, Margaryan A, Schroeder H, Orlando L, Willerslev E, Allentoft ME. 2015. Improving access to endogenous DNA in ancient bones and teeth. *Sci Rep* 5:11184.

Drummond AJ, Rambaut A. 2007. BEAST: Bayesian evolutionary analysis by sampling trees. *BMC Evol Biol* 7:214.

Ferreira MS, Jones MR, Callahan CM, Farelo L, Tolesa Z, Suchentrunk F, Boursot P, Mills LS, Alves PC, Good JM, et al. 2021. The Legacy of Recurrent Introgression during the Radiation of Hares.Buckley T, editor. *Systematic Biology* 70:593–607.

George D, Bizzarri C, Bianco P, Trentacoste A, Whitlam J, Best J. 2017. Recent Research in Cavità 254 (Orvieto, Italy). *Etruscan Studies* 20.

Katoh K, Standley DM. 2013. MAFFT Multiple Sequence Alignment Software Version 7: Improvements in Performance and Usability. *Molecular Biology and Evolution* 30:772–780.

Li H, Durbin R. 2009. Fast and accurate short read alignment with Burrows–Wheeler transform. *Bioinformatics* 25:1754–1760.

Marques JP, Seixas FA, Farelo L, Callahan CM, Good JM, Montgomery WI, Reid N, Alves PC, Boursot P, Melo-Ferreira J. 2020. An Annotated Draft Genome of the Mountain Hare (Lepus timidus).Mank J, editor. *Genome Biology and Evolution* 12:3656–3662.

McKenna A, Hanna M, Banks E, Sivachenko A, Cibulskis K, Kernytsky A, Garimella K, Altshuler D, Gabriel S, Daly M, et al. 2010. The Genome Analysis Toolkit: A MapReduce framework for analyzing next-generation DNA sequencing data. Genome Res. 20:1297–1303.

Oliva A, Tobler R, Cooper A, Llamas B, Souilmi Y. 2021. Systematic benchmark of ancient DNA read mapping. *Briefings in Bioinformatics* 22:bbab076.

Peltzer A, Jäger G, Herbig A, Seitz A, Kniep C, Krause J, Nieselt K. 2016. EAGER: efficient ancient genome reconstruction. *Genome Biol* 17:60.

Rambaut A. 2018. FigTree.

Rambaut A, Drummond AJ, Xie D, Baele G, Suchard MA. 2018. Posterior Summarization in Bayesian Phylogenetics Using Tracer 1.7.Susko E, editor. Systematic Biology 67:901–904.

Schubert M, Lindgreen S, Orlando L. 2016. AdapterRemoval v2: rapid adapter trimming, identification, and read merging. *BMC Res Notes* 9:88.

Trentacoste A. 2021. In the belly of the earth: bones and the closing of subterranean space in central Italy. In: Deschler-Erb, Sabine, et al. Roman Animals in ritual and funerary contexts. Deutsches Archäologisches Institut. P. 217-236.
